# Investigating the spectral features of the brain meso-scale structure at rest

**DOI:** 10.1101/2020.05.26.114488

**Authors:** Riccardo Iandolo, Marianna Semprini, Diego Sona, Dante Mantini, Laura Avanzino, Michela Chiappalone

**Affiliations:** Rehab Technologies, Istituto Italiano di Tecnologia, 16163 Genova, Italy; Pattern Analysis & Computer Vision, Istituto Italiano di Tecnologia, 16152, Genova, Italy; Neuroinformatics Laboratory, Fondazione Bruno Kessler, 38123 Povo (Trento), Italy; Research Center for Motor Control and Neuroplasticity, KU Leuven, 3001 Leuven, Belgium; Brain Imaging and Neural Dynamics Research Group, IRCCS San Camillo Hospital, 30126 Venice, Italy; Department of Experimental Medicine, Section of Human Physiology, University of Genova, 16132 Genova, Italy; Ospedale Policlinico San Martino, IRCCS, 16132 Genova, Italy

**Keywords:** Community Detection, Frequency-specificity, Meso-scale, Network Neuroscience, Resting State

## Abstract

Recent studies provide novel insights into the meso-scale organization of the brain, highlighting the co-occurrence of different structures: classic assortative (modular), disassortative and core-periphery. However, the spectral properties of the brain meso-scale remain mostly unexplored. To fill this knowledge gap, we investigated how the meso-scale structure is organized across the frequency domain. We analyzed the resting state activity of healthy participants with source-localized high-density electroencephalography signals. Then, we inferred the community structure using weighted stochastic block-modelling to capture the landscape of meso-scale structures across the frequency domain. We found that meso-scale modalities were mixed over the frequency spectrum, with a core-periphery structure predominance. Nevertheless, we also highlighted a selective increase of disassortativity in the delta and theta bands, and of assortativity in the low gamma band (30-50 Hz). We further described other features of the meso-scale organization by identifying those brain regions which, at the same time, i) exhibited the highest degree of assortativity, disassortativity and core-peripheriness (i.e. participation), ii) were consistently assigned to the same community, irrespective from the granularity imposed by WSBM (i.e. granularity-invariance). We defined those brain areas as Participation and Granularity Invariant. In conclusion, we observed that the brain spontaneous activity shows frequency-specific meso-scale organization which may support spatially distributed and local information processing.

## 1 Introduction

Functional connectivity (FC), i.e. the statistical association among neural signals of separate brain regions (Friston, 2011), has received a great deal of attention during the last years (Reid et al., 2019). FC has been widely recognized as a tool to investigate spatio-temporal properties of brain networks. These networks have been characterized at different levels of topological organization (Betzel & Bassett, 2017), ranging from local (single brain area or node) to global (whole-brain network) (Hallquist & Hillary, 2019), through the intermediate level referred to as meso-scale (Betzel, Medaglia, & Bassett, 2018). The single unit of the meso-scale architecture is a “community” (or module), which is composed by a set of nodes sharing similar connectivity patterns. Modules are crucial elements of FC network organization since they are essential to identify areas belonging to the same functional domain. Moreover, modules well describe network resilience and flexibility in response to external perturbation (as in the case of occurred cerebral lesions) and also they shape the information flow (Sporns, 2018). To date, the meso-scale structure of the human brain has been extensively investigated by community detection algorithms prone to detect “assortative” (also defined as “modular”) meso-scale structure (Betzel, Medaglia, et al., 2018; Newman, 2006), for a review see (Garcia, Ashourvan, Muldoon, Vettel, & Bassett, 2018). Briefly, in the assortative structure, the within-community densities are greater than the between-community densities. In other words, this structure facilitates information processing of segregated modules while the integration capability between them is reduced (Betzel, Bertolero, & Bassett, 2018). Non-assortative community interactions have been also described, such as the “disassortative” and the “core-periphery” (Betzel, Medaglia, et al., 2018). A disassortative structure is complementary to the assortative one. It is characterized by the connections between communities being greater than within communities, thus suggesting a strong flow of information between different modules. In the core-periphery structure, the nodes of a high-density *core* strongly interact with nodes of other *periphery* communities, which are characterized by poorly connected nodes. This structure allows an efficient broadcasting of information between core and peripheries (Betzel, Bertolero, et al., 2018). Importantly, it has been recently shown that these three classes (i.e. assortative, disassortative and core-periphery) also referred to as “community motif”, may coexist in the brain, forming the so-called mixed meso-scale structure (Betzel, Medaglia, et al., 2018; Garcia et al., 2018). Therefore, it is pivotal to detect the richness and diversity of meso-scale organization, without being limited by the analysis of assortative class only, as recently proposed (Betzel, Bertolero, et al., 2018; Betzel, Medaglia, et al., 2018). To this end, some algorithms have been presented in the literature (Fortunato, 2010), such as the Weighted Stochastic Block Model - WSBM (Aicher, Jacobs, & Clauset, 2014) able to capture the meso-scale diversity. An important feature of WSBM is the exploitation of the stochastic equivalence principle, according to which the network nodes belonging to a given community have the same probability of being connected with all the remaining nodes of the network (Aicher et al., 2014). The WSBM can thus detect other modalities of meso-scale modules interactions, beyond assortativity (Betzel, Medaglia, et al., 2018). Recent studies investigating human (Betzel, Bertolero, et al., 2018; Betzel, Medaglia, et al., 2018; Faskowitz, Yan, Zuo, & Sporns, 2018) and non-human brain networks (Faskowitz & Sporns, 2019; Pavlovic, Vertes, Bullmore, Schafer, & Nichols, 2014) made use of the WSBM method. In these investigations, human brain networks were derived with magnetic resonance imaging (MRI), using either functional (during both rest (Betzel, Medaglia, et al., 2018) and task (Betzel, Bertolero, et al., 2018)) or structural data (Betzel, Medaglia, et al., 2018; Faskowitz et al., 2018). Overall, the above results reported that assortativity dominates resting state FC with the co-existence of non-assortative communities (Betzel, Bertolero, et al., 2018), thus indicating that brain networks are not characterized by a unique community structure. Nevertheless, how the resting state FC meso-scale structure is organized across the frequency domain still needs to be investigated. Thus, to fill this knowledge gap, we explored the spectral properties of the brain meso-scale. To reach this goal, we exploited high-density electroencephalography (hdEEG), a non-invasive electrophysiological technique. Notably, hdEEG provides a unique opportunity to capture the richness of neuronal oscillations’ spectral content (Siegel, Donner, & Engel, 2012), and it was recently employed to reconstruct and unravel novel features of human brain activity during resting state in health (Liu, Farahibozorg, Porcaro, Wenderoth, & Mantini, 2017; Samogin, Liu, Marino, Wenderoth, & Mantini, 2019; Seeber et al., 2019) and disease (Cassani, Estarellas, San-Martin, Fraga, & Falk, 2018; Coito et al., 2016; Damborská et al., 2020; Waninger et al., 2020). By coupling hdEEG recordings with appropriately built head model conductors and with source reconstruction algorithms, it is possible to achieve neural sources reconstruction with relatively good (i.e. in the order of less than 1 cm) spatial resolution (He, Sohrabpour, Brown, & Liu, 2018; Seeber et al., 2019). This permitted the estimation of large-scale resting state networks that spatially overlap with those obtained with functional MRI (fMRI) (Liu et al., 2017) and magnetoencephalography (MEG) (Coquelet et al., 2020). Thus, we posit that describing the spectral features of FC meso-scale architecture estimated from source-localized hdEEG recordings will have important implications to highlight novel properties of the human brain at rest (de Pasquale, Corbetta, Betti, & Della Penna, 2018; Hipp, Hawellek, Corbetta, Siegel, & Engel, 2012; Siems, Pape, Hipp, & Siegel, 2016).

With this aim, we here exploited the peculiar features of hdEEG-based source imaging, to identify modules of spontaneous oscillatory activity and to characterize their properties. Specifically, we tested whether the meso-scale structure is frequency-dependent, by examining if assortative, disassortative and core-periphery community motifs are tuned onto a specific frequency or if they are equally distributed over the frequency domain. To address these questions, we applied the WSBM to FC adjacency matrices estimated from source-localized hdEEG recordings of healthy participants (Iandolo et al., 2020; Liu et al., 2017; Samogin et al., 2019), in order to define the spatial cerebral distribution of modules at different number of partitions (K^th^). Then, we described the assortative, disassortative and core-periphery community interactions across the frequency spectrum and we identified those brain regions exhibiting the highest degree of each community motif. Interestingly, we also observed that some regions were consistently assigned to the same community, irrespective from the granularity (i.e. the K partitions) that we imposed to the WSBM algorithm. We then identified brain areas which were, at the same time, maintained across partitions and exhibited the highest degree of assortativity, disassortativity and core-peripheriness. We defined those areas as Participation and Granularity Invariant.

Overall, we observed that the spontaneous oscillatory activity relies on frequency-specific topological meso-scale organization, which may support spatially distributed and local information processing.

## 2 Materials and Methods

### 2.1 Participants

We recruited 29 healthy volunteers (28.8 ± 3.6 years, mean ± SD, 14 females). To be included, the participants had: *a)* to be right-handed according to the Edinburgh inventory (Oldfield, 1971); *b)* to be without neurological or psychiatric disorders; *c)* to have normal or corrected-to-normal vision; *d)* to be free of psychotropic and/or vasoactive medication. Prior to the experimental procedure, all participants provided written informed consent. The study, which was in line with the standard of the Declaration of Helsinki, was approved by the local ethical committee (CER Liguria Ref. 1293 of September 12^th^, 2018).

### 2.2 Resting state hdEEG recording and MRI acquisition

HdEEG signals were recorded using a 128-channel amplifier (ActiCHamp, Brain Products, Germany) while participants were comfortably sitting with their eyes open fixating a white cross on a black screen for five minutes. Participants were required to relax as much as possible and to fixate the cross, located in the middle of a screen in front of them. The experiment was performed according to the approved guidelines, in a quiet laboratory with soft natural light. HdEEG signals were collected at 1000 Hz sampling frequency, using the electrode FCz (over the vertex) as physical reference electrode. The horizontal and vertical electrooculograms (EOG) were collected from the right eye for further identification and removal of ocular-related artifacts. Prior to resting state hdEEG recordings, the three-dimensional locations of the 128 electrodes on the scalp were collected with either infrared color-enhanced 3D scanner (Taberna, Guarnieri, & Mantini, 2019) or Xensor digitizier (ANT Neuro, The Netherlands). To build each participant’s high-resolution head model, the participants underwent T1-weighted MRI acquisition using either a 3 T (N = 25) or a 1.5 T (N = 4) scanner (see Suppl. Materials for details about T1-weighted images acquisition parameters).

### 2.3 Pre-processing of hdEEG recordings

HdEEG preprocessing was performed according to the same steps described in previous works (Liu et al., 2017; Samogin et al., 2019). Briefly, we first attenuated the power noise in the EEG channels by using a notch filter centered at 50 Hz. Later, we identified channels with low signal to noise ratio by following an automatic procedure. We combined information from two channel-specific parameters: i) the minimum Pearson correlation between a channel against all the others in the frequency band of interest (i.e. 1-80 Hz); ii) the noise variance that we defined in a band where the EEG information is negligible (i.e. 200-250 Hz). We defined a channel as “bad”, whenever one of the two parameters described above were outliers as compared to the total distribution of values. We interpolated the identified bad channels with the information of the neighboring channels, using Field Trip (http://www.fieldtriptoolbox.org/). Then, hdEEG signals were band pass filtered (1-80 Hz) with a zero-phase distortion FIR filter and downsampled to 250 Hz. To further reduce noise in our data, we employed the fast-ICA algorithm (http://research.ics.aalto.fi/ica/fastica/) to identify independent components related to ocular and movement artifacts. To classify the ocular artifacts we used the following parameters: i) Pearson correlation between the time-course of the independent components and the vertical and horizontal EOG; ii) the coefficient of determination obtained by fitting the independent component (IC) spectrum with a 1/f function. We classified the IC as ocular artifacts if at least one of the two parameters was above a pre-defined thresholds (0.2 and 0.5, as in (Liu et al., 2017)). Finally, for movement-related artifacts, we used the kurtosis of the independent component (we considered a kurtosis exceeding the value of 20 (Liu et al., 2017) indicative of a noisy IC). We re-referenced the artifacts-free signals with the average reference approach (Liu et al., 2015).

### 2.4 Head model of volume conduction and source reconstruction

We followed the same procedure as detailed in (Iandolo et al., 2020). Briefly, we used T1-weighted structural images in order to generate a realistic volume conductor model. In accordance with previous studies (Liu et al., 2017; Samogin et al., 2019), we assigned conductivity values to 12 tissue classes (skin, eyes, muscle, fat, spongy bone, compact bone, gray matter, cerebellar gray matter, white matter, cerebellar white matter, cerebrospinal fluid and brainstem), based on the literature (see (Liu et al., 2017) for the conductivity values assigned per each tissue class). Then, given the intrinsic difficulty in segmenting all the 12 classes directly on the T1-weighted individual space, we warped the MNI (Montreal Neurological Institute) template to individual space using the normalization tool of SPM12 (http://www.fil.ion.ucl.ac.uk/spm/software/spm12), as reported in (Liu et al., 2017). Then, we spatially co-registered the 128 electrodes positions onto each individual T1-weighted space. We approximated the volume conduction model using a finite element method (FEM) and we employed the Simbio FEM method (https://www.mrt.uni-jena.de/simbio/) to estimate the relationship between the measured scalp potentials and the dipoles corresponding to brain sources. Finally, by combining the individual head model conductor and the artifacts-free hdEEG signals, we reconstructed source activity using the eLORETA (Pascual-Marqui et al., 2011) algorithm. Sources were constrained within a 6 mm regular grid covering the gray matter as in (Liu et al., 2017). Thus, we reconstructed the sources (voxels) per each participant and we then mapped the voxels time-courses into 384 regions of interest (ROIs) of the AICHA atlas (Joliot et al., 2015). To this purpose, we estimated the activity of each ROI employing the first principal component of the voxels falling within a sphere centered in the ROI center of mass and with 6 mm radius. This procedure defines the nodes for the subsequent FC meso-scale structure investigation.

### 2.5 Spectral analysis and definition of connectivity matrix

We decomposed each ROI time-course using Short-Time Fourier Transform. We used a frequency bin of 1 Hz in the range [1÷80 Hz] and a Hamming window of 2 seconds duration with 1 second overlap between consecutive windows. Then, for each participant, we estimated the FC adjacency matrices per each frequency bin (80 in total) by calculating the pairwise connection strength among the ROIs. To estimate connection strength we employed the method of power envelope orthogonalization (Hipp et al., 2012) that estimated the amplitude-amplitude coupling. Indeed, although the brain activity estimation at the sources level is a promising tool to investigate the brain dynamics at both good spatial and high temporal resolutions, it is affected by the signal leakage problem (Hipp et al., 2012; Siems et al., 2016). Reconstructing cortical and sub-cortical sources (several thousand sources) from scalp potentials (here 128 electrodes) is an ill-posed inverse problem, introducing artefactual cross-correlations between sources. A recent validation study (Siems & Siegel, 2020) established the power envelope orthogonalization as an effective candidate to estimate the physiological FC properties in the field of neuroimaging by electrophysiological recordings. Thus, we followed the same orthogonalization procedure described and employed in previous EEG studies (Samogin et al., 2019; Siems et al., 2016). Then, per each participant, we averaged the adjacency matrices (i.e. see above) within the following frequency ranges, according to (Samogin et al., 2019; Zhao, Marino, Samogin, Swinnen, & Mantini, 2019): delta (δ, 1-4 Hz), theta (θ, 4-8 Hz), alpha (α, 8-13 Hz), beta (β, 13-30 Hz), gamma low (γ_L_, 30-50 Hz) and gamma high (γ_H_, 50-80 Hz). By averaging the subjects’ connectivity matrices *ADJ*_*S*_, we obtained the group level adjacency matrices per each frequency band 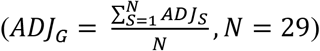. Then, we linearly mapped each *ADJ*_*G*_edges weights between the interval [0, +1]:

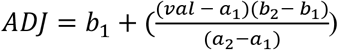

where *val* is a single element of *ADJ*_*G*_; *a*_1_, *a*_2_ are the minimum and maximum edges value of *ADJ*_*G*_; *b*_1_, *b*_2_ are the limits of the new range 0 and 1. This transformation allows for further comparison of the meso-scale structure among different frequency content. It is indeed necessary to normalize the weights of the adjacency matrices in the same range to compare outputs of the WSBM, according to the literature (Aicher et al., 2014). The single-subject level adjacency matrices (*ADJ*_*S*_) followed the same mapping procedure so that edge weights would fall in the interval [0, +1].

### 2.6 Community detection via Weighted Stochastic Block Models

WSBM is as an unsupervised learning algorithm for the identification of network communities that group together network nodes that have similar FC patterns (Aicher et al., 2014). The WSBM can work without the need of thresholding the adjacency matrix, as this procedure might have a negative impact on the properties of the inferred community structure, as previously reported (Aicher et al., 2014). The WSBM goal is to learn the hidden community structure that is estimated from both the existence and the weights of edges. Moreover, the algorithm retains the principle of stochastic equivalence, which differentiates this community detection approach from the modularity maximization algorithms, employed for community detection in network neuroscience but biased towards assortativity (Betzel, Bertolero, et al., 2018; Betzel, Medaglia, et al., 2018). Additionally, it is important to note that stochastic block-modelling has the unmet advantage of being a generative model, as it tries to estimate the process underlying the observed network topology. The WSBM learns two parameters starting from the adjacency matrix and from a priori assumptions about the distributions of weights and existence of edges. An important parameter is the vector of nodes assignment *Z* = [*z*_1_, …, *z*_*N*_] where *z*_*j*_ *ε* {1, . ., *K*}, with N the number of nodes and K the number of communities the algorithm must learn. The other parameter is the edge bundle matrix (or affinity matrix) θ ([*K* × *K*]), representing the probability of two communities being connected. It is worth noting that the probability of connection between two nodes only depends on their community labels assignment, 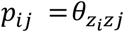. In its formulation, the log-likelihood of the adjacency matrix being described by the parameters θ and *Z*, can be written as (Aicher et al., 2014; Betzel, Medaglia, et al., 2018):

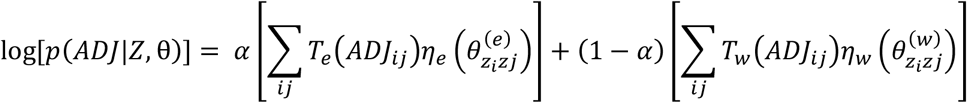

where α is a tuning parameter that combines the contribution of the two summations, which respectively model edges weights and edges existence, to infer the latent community structure. 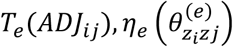 and 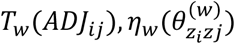 are the sufficient statistics and the natural parameters of the exponential family describing the distributions of the edges existence (*T*_*e*_, *η*_*e*_) and the edges weights (*T*_*w*_, *η*_*w*_). Lastly, *i, j* indicate the edges of the adjacency matrix onto which we inferred the latent community structure. Usually, when applying the WSBM framework to structural and functional brain networks, the edges existence and weights are drawn from Bernoulli and Normal distributions (Aicher et al., 2014; Betzel, Bertolero, et al., 2018; Betzel, Medaglia, et al., 2018), respectively. In our case, α is set to zero because the graph is fully connected (i.e. no thresholding applied) and thus we did not need to model the edges existence. Hence, our likelihood maximization is simplified leading to a pure-WSBM (Aicher et al., 2014) (WSBM indicates pure-WSBM throughout the text) that learns from the weights information, that are assumed to be normally-distributed between communities. The remaining issue is to find a reliable estimation of the posterior distribution, i.e. *p*(*Z, θ*|*ADJ*) that has no explicit analytic formulation (Aicher et al., 2014). To this purpose, we made use of the code freely available here (http://tuvalu.santafe.edu/~aaronc/wsbm/). The code finds an approximation of the posterior probability using a Variational Bayes (VB) approach. VB provides a solution to approximate the unknown posterior distribution by transforming an inference problem into an optimization problem. The algorithm minimizes the Kullback-Lieber divergence D_KL_ (Fox & Roberts, 2012) to the posterior probability (for further information about D_KL_ applied to WSBM, see (Aicher et al., 2014)). The solution proposed by (Aicher et al., 2014) states that minimizing the D_KL_ is equivalent to maximize the evidence lower bound of the model marginal log-likelihood (logEvidence), *p*(*ADJ*|*Z, θ*). Thus, the best approximation of the posterior is obtained through a procedure aimed at maximizing the logEvidence score. If the logEvidence is maximized, the D_KL_ is the closest possible to the posterior distribution, *p*(*Z, θ*|*ADJ*). After properly initializing the priors for *θ* and *z*, the VB algorithm takes the best (i.e. the greatest) logEvidence value across multiple independent trials (or restarts) of the algorithm. We choose a maximum of 100 independent trials to find the best logEvidence value. Within this limit, the algorithm searches for the best logEvidence value. At each trial, the initial probability of a node being assigned to a community is randomized. Every time a better logEvidence value (i.e. a better solution) is obtained, the algorithm updates the solution. We selected the communities assignment with the greatest logEvidence value. We run the WSBM model for different values of K (ranging from 3 to 8) and, we performed 100 WSBM fits per each value of K.

### 2.7 Community assignment

In order to choose the best nodes assignment among all the WSBM fits, we used the community assignment corresponding to the central fit across the WSBM fits. We defined the central fit as the fit whose distance is minimized from all the others fits using the Normalized Variation of Information (NVI), as in a previous work (Faskowitz et al., 2018) (we used the function *partition_distance*.*m* of the Brain Connectivity Toolbox (Rubinov & Sporns, 2010)). We used the central fit not only to identify and to show the resulting communities at the group level, but also for all the subsequent steps of our analysis: the investigation of how the total amount of between-community interactions varies across frequency bands. Indeed, in addition to fit the WSBM generative model to *ADJ*, we also fitted the model for all K values (i.e. from 3 to 8) at the single-subject level (*ADJ*_*S*_). Thus, for each participant, we performed 100 WSBM fits and we selected as best fit the central one according to the NVI, as for the group level. The central fit was calculated for each of the six frequency bands.

### 2.8 Characterizing the meso-scale structure: community motifs

At the single-subject level, we investigated, for each choice of K, how pairs of communities interacted with each other in order to generate assortative, disassortative and core-periphery architecture. This permitted us to investigate the spectral organization of community motifs. For each pair of communities *r* and *c*, we estimated the within- and between-community density (Betzel, Bertolero, et al., 2018), a topological property of the detected modules (Garcia et al., 2018):

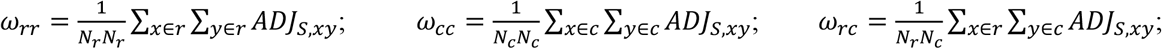

Where, *N*_*r*_ and *N*_*c*_ are the number of nodes assigned to the communities *r* and *c* at the central fit. We calculated community density for the different bands at a given K partition. Then, the between-community motifs (*M*_*rc*_) fall into one of the three categories as reported in (Betzel, Bertolero, et al., 2018; Betzel, Medaglia, et al., 2018), according to the following criteria:

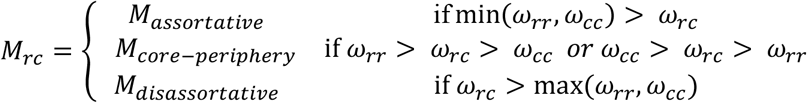

We calculated the percentage of between-community interactions for each participant with respect to the total number of possible interactions, corresponding to 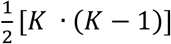. Then, we averaged the occurrence of each community class across all the different K partitions (K = 3…8) to identify a measure which can globally consider the entire range of meso-scale granularity: we referred to it as “global” community motif modality.

### 2.9 Statistical analysis of motif interaction

To answer the question of the frequency-specificity of between community interactions, we needed to identify whether and how global motif modality changes across frequency bands. Given the non-normality of the motifs distributions, we performed a set of Kruskal-Wallis non-parametric tests to examine, for each community motif class (i.e. assortative, disassortative and core-periphery), whether the frequency band has a statistically significant effect. Then, we employed a post-hoc comparison of mean ranks as implemented in Statistica 13 software package (Statsoft Inc., Tulsa) to further highlight the potential differences among the six bands within each community class. Furthermore, we aimed to identify those brain areas which presented the greatest amount of assortativity, disassortativity and core-peripheriness (three independent tests). To achieve this goal, we built each motifs’ null distribution by randomly shuffling each subject node-level motifs (number of permutations: 10^4^). Then, we obtained a mean null distributions by averaging across participants. We computed the 99^th^ percentile of this null distribution and we searched for those nodes belonging to the empirical distribution whose motifs’ values fell above this value (p < 0.01). Finally, we labelled these significant areas with the same name assigned by the AICHA atlas.

### 2.10 Identifying invariant behavior of communities and nodes across K partitions

We also investigated how meso-scale organization changes for different K partitions and explored the presence of nodes always clustered together regardless of the specific K choice. For each frequency band we proceeded as follows. For each choice of K, ranging from 3 to 8, we first calculated how many nodes of each cluster of the K-th partition fell in each cluster of the (K+1)-th partition. We were therefore able to observe how communities obtained in the K-th partition, diverge (branch) and/or converge (flow) into the communities obtained in the (K+1)-th partition. Then, we investigated whether and how some nodes of the three clusters obtained for K = 3 were maintained together in the partitions obtained for K > 3. We defined such nodes as invariants to the WSBM-clustering procedure and identified them as those nodes meeting the following requirements: (*i*) nodes belonging to the largest portion of the cluster found in the K-th partition; and (*ii*) nodes belonging to the greatest flow reaching the cluster found in the (K+1)-th partition. Finally, we aimed at identifying those nodes that were simultaneously invariant to the K-th partition and also significantly exhibiting one specific community motif. We defines these nodes as Participation and Granularity Invariant (PGI nodes throughout the text).

## 3 Results

In this study, we reconstructed neural sources per each participant and we then mapped them onto 384 ROIs of the AICHA atlas (Joliot et al., 2015). This procedure defined the nodes for the subsequent meso-scale structure investigation. We then extracted the FC adjacency matrices using power envelope orthogonalization and we applied the WSBM (for different K values, ranging from 3 until 8). We investigated the organization of the meso-scale structure across canonical frequency bands: delta (δ, 1-4 Hz), theta (θ, 4-8 Hz), alpha (α, 8-13 Hz), beta (β, 13-30 Hz), gamma low (γ_L_, 30-50 Hz) and gamma high (γ_H_, 50-80 Hz). We then described community motif organization across frequency bands, and we identified those brain areas which i) showed the highest community motifs’ degree (i.e. statistically significant participation); ii) were consistently assigned to same community for increasing meso-scale granularity (i.e. granularity invariant); iii) addressed, at the same time, the two previous criteria (i.e. significant participation and granularity invariance - PGI nodes).

### 3.1 Spectral analysis of meso-scale connectivity structure

We examined the spatial distribution of the community assignments (K = 3…8) across frequency bands. In general, clusters are less sparse and more compact when moving from low towards high rhythms (see Figure 1 and Supplementary Figures 1-5). Let us take, as a representative example, the spatial distribution of five communities (K = 5, Figure 1), which lies in the middle of the selected K range. First, it resembles the pattern observed for all partitions: increasing the frequency band corresponds to a compact and mirror symmetric (with respect to brain midline) representation of clusters (see also Supplementary Materials Figures 1-5). As for the spatial distribution of the communities in the δ band (see Figure 1, panel a1), we obtained an association cluster, mostly located in the right hemisphere (corresponding roughly to somatic areas, and association parieto-temporo-occipital (PTO) cortex, red). Another lateralized cluster was obtained in the left hemisphere, putatively associated with executive functions (frontal and temporal lobe, purple). We found a sparse bilateral cluster, predominantly located in the right hemisphere PTO cortex (yellow). Finally, other two bilateral fragmented clusters were spanning several areas (prefrontal areas including primary and premotor cortices as well as parietal and temporal lobes, blue and green). They presented high-level of sparseness in laterally located regions while they were consistent in right medial regions (blue) and left medial regions (green).

**Fig. 1.**
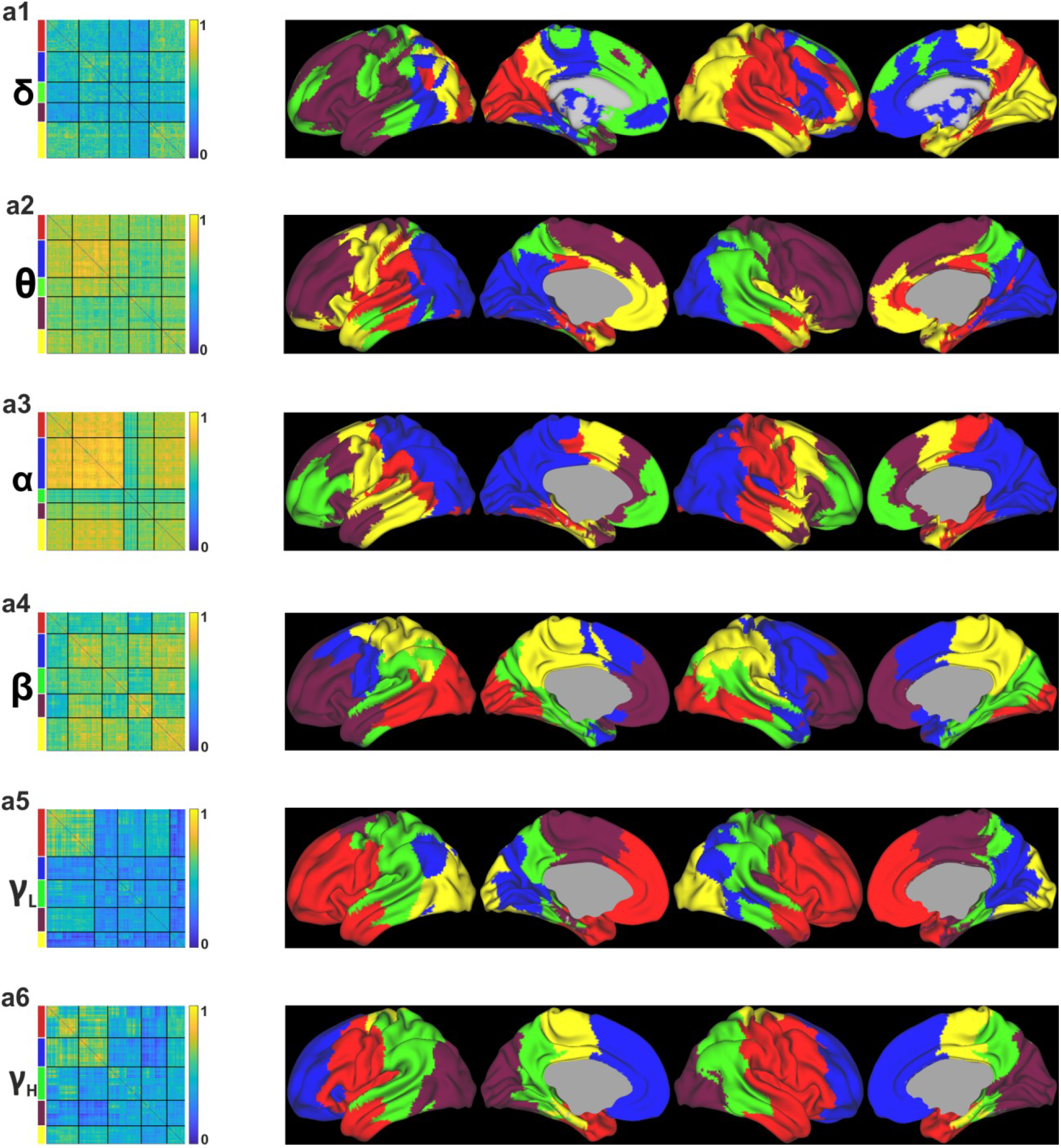
Organization of the meso-scale structure in the frequency domain. Each row represents the K = 5 community assignments in each of the considered frequency band: δ (**a1**), θ (**a2**), α (**a3**), β (**a4**), γ_L_ (**a5**), γ_H_ (**a6**). Each row contains the re-ordered group level adjacency matrix after WSBM estimation and spatial distribution of the five partitions across the brain. Colors on left side of each adjacency matrix match with the colors overlaid on the brain.

As for the θ oscillations (see Figure 1, panel a2), the block-modelling partitioning associated brain areas bilaterally in the frontal lobes (purple), and in two bilateral PTO clusters: one more posterior (blue) than the other (green). Moreover, the PTO-anterior (green) mainly covered the right hemisphere whereas PTO-posterior resembled a more symmetric pattern between hemispheres. The remaining two clusters were scattered bilaterally across the frontal lobe (yellow), and in the cingulate gyrus and left parieto-temporal cortex (red).

As for the α rhythm (see Figure 1, panel a3), the generated cluster approximately followed symmetric patterns across hemispheres: partitions were the bilateral prefrontal cortex (green), premotor cortices and temporal lobes (purple), primary motor and temporal cortices (yellow), right temporo-parietal cortex (red) which showed a less bilateral pattern than previous partitions as it localized predominantly around the lateral sulcus in the left hemisphere. The latter cluster belonged to PTO cortex bilaterally (blue) roughly resembling the “dorsal/where stream” originating from the occipital areas. Moreover, it covered most nodes which are contained in the PTO-posterior cluster of the previous θ band.

As for the β band (see Figure 1, panel a4), the clusters covered bilaterally prefrontal cortices and a small portion of the left temporal lobe (purple), motor and premotor areas (blue), parietal areas (yellow), PTO cortices (both red and green clusters). However, the more posterior-PTO partition spanned the primary and high-order visual areas as well as a portion of the temporal areas which approximately resembles the “ventral/what stream”, while the more anterior-PTO cluster did not include the primary visual cortex.

As for the γ_L_ oscillations (see Figure 1, panel a5), the clustering showed a bilateral frontal partition that mainly gathered prefrontal and premotor cortices (red). This cluster roughly merged together the two separate frontal clusters of the β rhythm (blue and purple). Other partitioning corresponded to frontal and parietal cortices (purple), PTO association cortices (green and blue) and occipital areas (yellow).

Finally, as for the high γ_H_ (see Figure 1, panel a6), we found a bilateral frontal cluster (blue), bilateral frontal, parietal and temporal (red), parietal cluster more located in medial regions (yellow), bilateral PTO (green), occipital (purple). This last cluster corresponds to the occipital cluster of the γ_L_.

Overall, for brain areas close to the midline, we found more compact and symmetric spatial distribution of communities than in the laterally-placed areas. Indeed, when moving towards more lateral regions, the clusters spatial pattern became more complex than the one observed in medial areas. Lower rhythms (δ, θ) presented more shattered clusters than mid-low (α, β) and high oscillations (γ_L_, γ_H_). Indeed, we found that higher rhythms were more likely characterized by distinct and less fragmented clusters than lower frequency bands, reflecting the general behavioral of the other partitioning (i.e. K = 3, 4, 6, 7, 8, see Supplementary Materials, Figure 1-5).

### 3.2 Non-assortativity of community structure in the frequency domain

To investigate whether meso-scale structure is frequency-specific, we evaluated possible differences among the six frequency bands considering all three community classes (i.e. assortative, disassortative, core-periphery) for each choice of the K parameter. We then averaged the occurrence of each community class across different partitions obtained for different values of K (K = 3…8). We found a statistically significant effect of the frequency band for the assortative (p = 0.0024) and disassortative (p < 0.0001) structure, as revealed by Kruskal-Wallis non-parametric testing (see Figure S6 for an overview of motifs community interaction at each considered K-value). Specifically, when considering the assortative structure the γ_L_ rhythm showed significant increase with respect to the δ and θ rhythms, see Figure 2, panel c. On the other hand, modules of spontaneous activity interacted more disassortatively in the δ and θ bands than the γ_L_ and γ_H_ bands, see Figure 2, panel d. By contrast, for the core-periphery class, we did not find any significant difference across the six bands (p = 0.11), suggesting that the core-periphery structure is homogeneously distributed across frequency bands (see Figure 2, panel a). Albeit not statistically significant, core-peripheriness was the most predominant community motif across subjects and partitions (see also Figure S6).

**Fig. 2.**
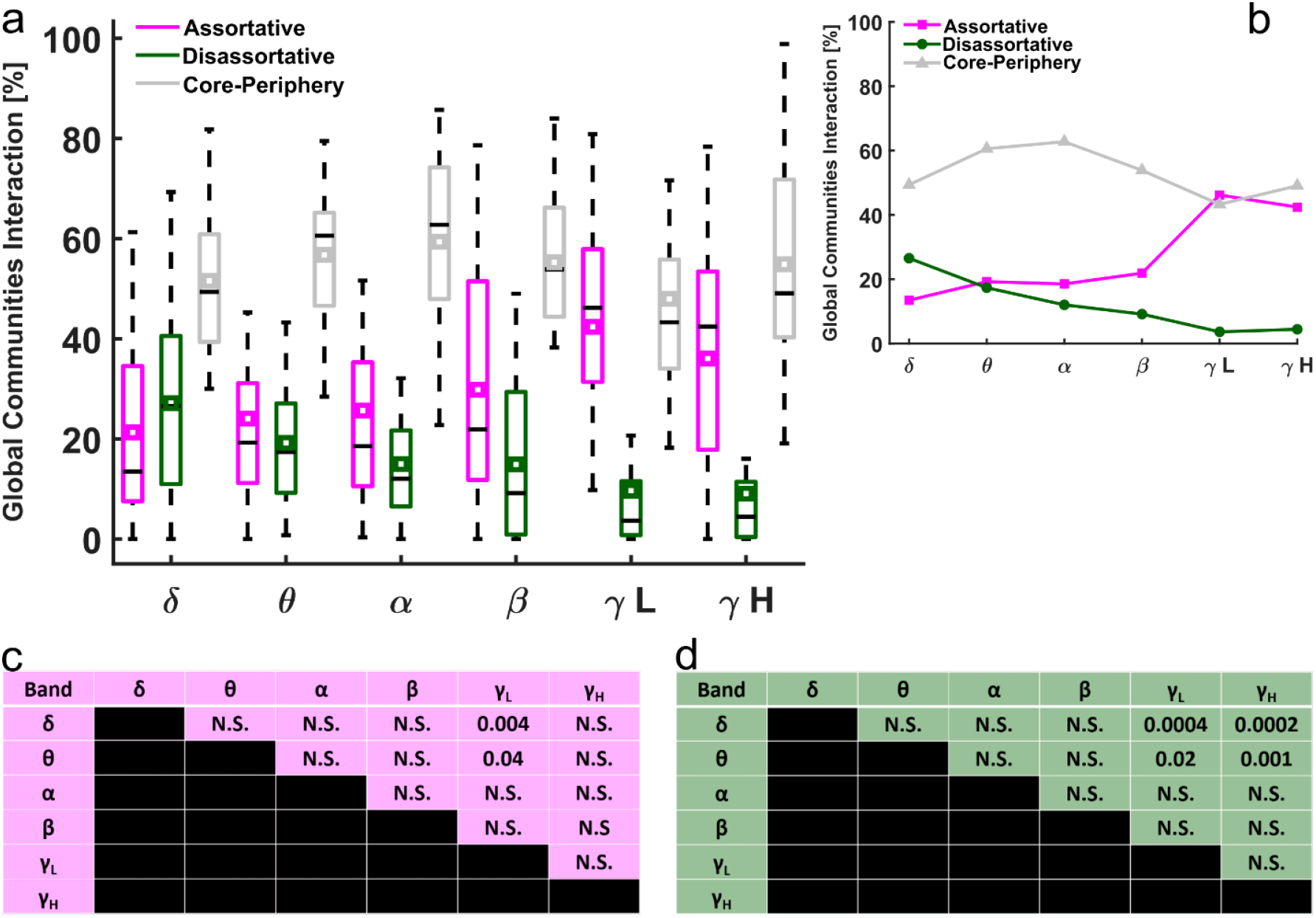
Organization of the meso-scale structure in the frequency domain. a Boxplots representing distributions across participants of the three meso-scale motifs for each frequency band. Magenta: assortative; green: disassortative; gray: core-periphery. Boxplots show upper and lower bound of the distributions at 25^th^ and 75^th^ percentile. Whiskers extend to the most extreme data points not considered outliers. The black horizontal lines represent the median, while the small colored squares indicate the mean of the distributions. **b** Median values of each meso-scale structure distributions (black horizontal lines in panel a) across frequency bands. **c-d**. Post-hoc comparison of mean ranks across frequencies. Tables highlight statistically significant assortative and disassortative between-communities interaction, panel c and d, respectively. Note that core-periphery interactions across bands were non-statistically significant and thus we did not perform the multiple comparison test.

Overall, we observed, within the assortative and disassortative classes, complementary trends along the entire range of oscillatory rhythms (with significant difference between some low and high bands, i.e. δ and θ vs. γ_L_). Specifically, for increasing frequencies we found a decreasing disassortative and an increasing assortative trend, respectively (see Figure 2, panel b).

In order to appreciate motifs distribution at the node level, we averaged across participants the total amount of meso-scale modalities obtained in each node for different choices of K (K = 3…8) and we then overlaid these values (at the node level) onto the T1-weighted template (see Figure 3). We observed that the core-periphery structure was clearly predominant in all frequency bands and for most brain areas. Instead, assortative and disassortative modalities exhibited trends across bands and whole-brain variations within the same band. The PTO areas in γ_L_ showed the highest degree of assortativity (see Figure 3, panel a). In this band, there was a spatial gradient in assortativity increasing from anterior to posterior areas. Disassortative structure peaked in δ and θ bands, with a whole-brain maximum in δ (see Figure 3, panel a). We performed a custom non-parametric test to identify areas with the highest degree of assortativity, disassortativity and core-peripheriness (see Figure 3, panel b). As for the assortative class, significant areas were located in temporal cortex, precuneus and nucleus caudate for the δ band (subcortical nuclei are not overlaid in Figure 3, see Table S2). Frontal, superior and middle frontal areas for α (bilaterally) and β (left hemisphere) band, while bilateral areas of the PTO cortex for γ_L_ and γ_H_ band (see Supplementary Materials, Tables S2-S7 for further information). As for the disassortative structure, significant areas were mostly located in temporal gyrus, insula and in subcortical areas such as caudate nucleus, putamen and thalamus for δ band, whereas thalamus and occipital areas for θ band (again, subcortical areas are not depicted, for their description see Table S2 (for δ) S3 (for θ)). Precuneus, posterior cingulate cortex and parietal areas were instead significant for the β band. It is worth noting that thalamus was significantly disassortative in all bands (with the exception of γ_L_, see table S2-S7). Finally, the overall level of core-peripheriness was mostly uniform in the whole-brain, resulting in generally sparse and less significant brain areas than for the other two motifs. Notably, for slow oscillations we found significance mostly in the occipital and temporal areas (δ) as well as in the frontal, motor areas and insula (θ).

**Fig. 3.**
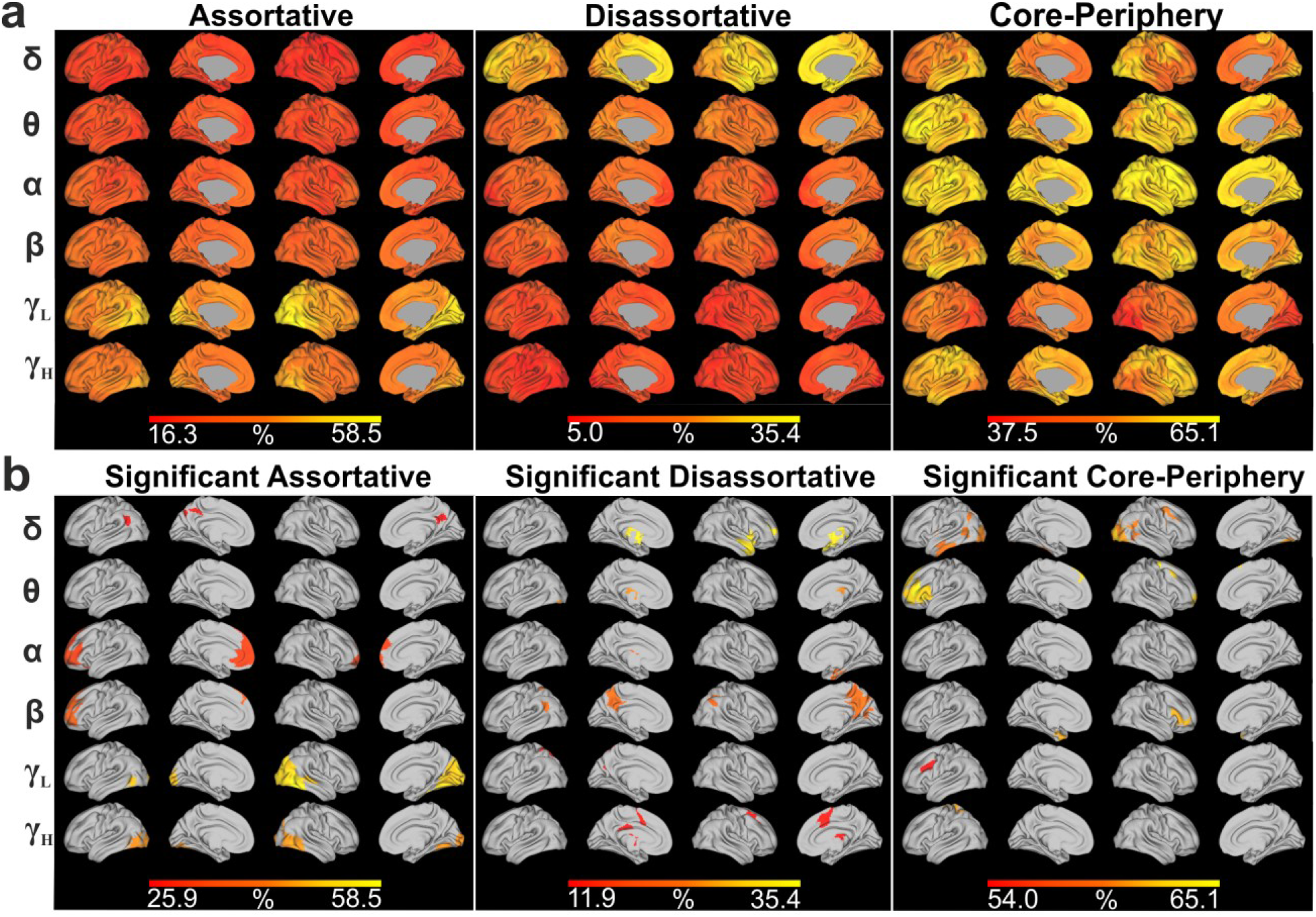
Global mean community interactions in the frequency domain across participants. a Each group of four columns indicates the meso-scale class (assortative, disassortative, core-periphery) while each row indicates the frequency band. The color-bar is customized between minimum and maximum values within each meso-scale modality. **b**. Significant nodes according to the community motifs, as revealed by the non-parametric permutation test. Columns and rows as in panel a. See also Supplementary Table S2, S3, S4, S5, S6, and S7 for further information about each significant node’s name and MNI coordinate, according to the definition given in the AICHA atlas.

### 3.3 Meso-scale invariants across partitions

We further characterized the meso-scale structure by looking at invariant modules across partitions. We observed how nodes belonging to the three communities in K = 3 were maintained clustered together or, on the contrary, they were assigned to other partitions from K > 3. With this procedure, we identified modules whose brain regions were assigned by the WSBM to the same clusters independently from the number of partitions required to be detected (see alluvial plots in Figure 4). Each transition identified the assignment of each clusters from K = 3 until K = 8 partitions. When nodes were clustered together across partitions, we assigned the same color-code to the corresponding flow (i.e. light-red, yellow and dark blue, see Figure 4). Albeit communities reconfigured across K-values, we highlighted invariant meso-scale pattern, independently from the K^th^ partition.

**Fig. 4.**
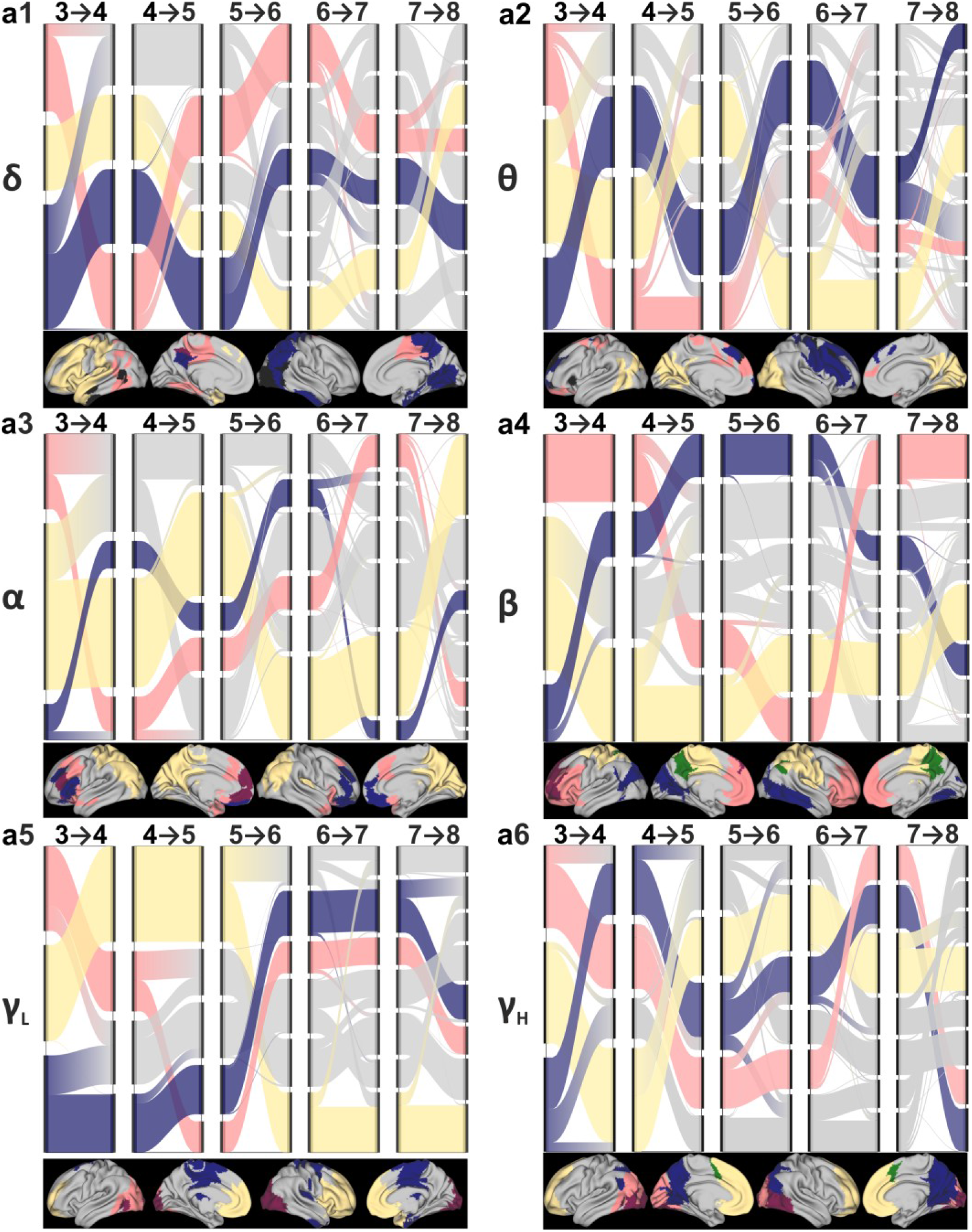
Communities reconfiguration across K-th partitions. Alluvial plots indicating 3 set of nodes (light-red, yellow and dark blue) assigned to the same community regardless of the partitions in each band -δ (**a1**), θ (**a2**), α (**a3**), β (**a4**), γ_L_ (**a5**), γ_H_ (**a6**). Gray flows indicate nodes failing to address the criteria defining main flows across partitions. Every time a colored flow’s branch fades towards gray indicates that nodes terminated in a different cluster. Vertical black lines on each side of the transitions indicates originating (from the lower-grain partition – left side) and arrival clusters (to the adjacent higher-grain partition – right side).

We further identified those brain areas that belonged to these invariant clusters and, at the same time, showed a maximally significant amount of between community interactions (see Figure 5 and Supplementary Tables S8-S13). Interestingly, different areas across bands addressed these two criteria, and we defined these areas as PGI. Several occipital and temporal areas which showed significant level of core-peripheriness (see Figure 3, panel b), were also found invariant to the number of communities imposed in the δ band (see Figures 5, panel a1 and Table S8). The same held for superior, middle frontal gyrus and precentral sulcus, but for the θ band (see Figures 5, panel a2 and table S9). Instead, we found that a large portion of left frontal areas in the α band had a significant level of assortativity (see Figure 3, panel b) and were assigned to same community across partitions (see Figures 5, panel a3 and Table S10). The β band, unlike the three previous rhythms, exhibited PGI areas in all three community motifs: left frontal areas (assortative), various parietal areas, cingulate and posterior cingulate cortex (PCC), precuneus and thalamus (disassortative), and right inferior frontal areas (core-periphery) (see Figures 5, panel a4 and Table S11). Both γ_L_ and γ_H_ exhibited PGI regions in occipital and temporal areas (assortative), and for γ_H_ only also in cingulate sulcus and supplementary motor area (disassortative) (See Figures 5, panel a5 and a6, and Table S12 and S13).

**Fig. 5.**
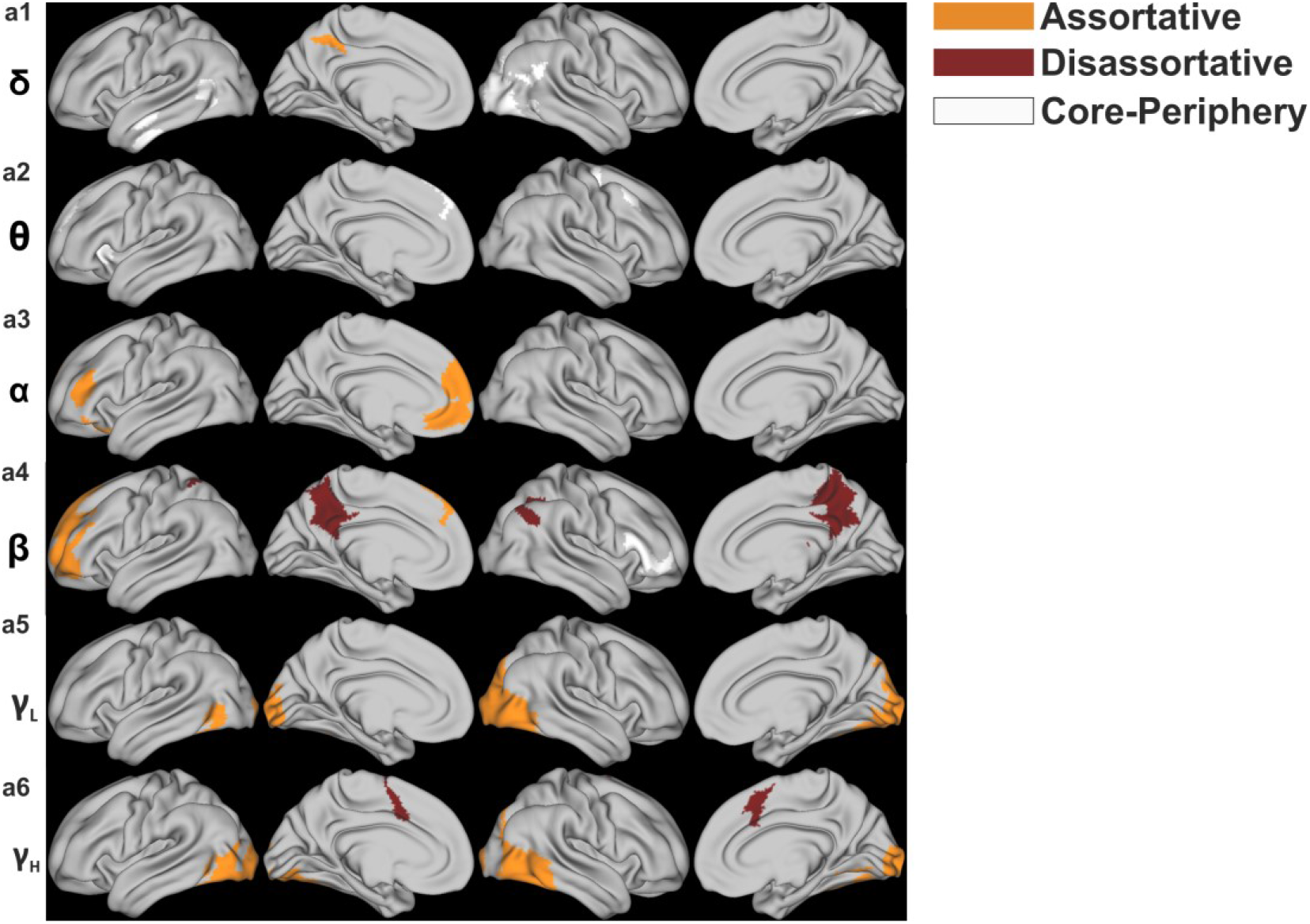
Participation and granularity invariant regions. Highlighted areas indicate areas that both belong to an invariant community across partitions and show significant level of assortativity (PGI-assortative areas, orange), disassortativity (PGI-disassortative, dark red) and core-peripheriness (white, PGI-core-periphery). Each row indicates a frequency band -δ (**a1**), θ (**a2**), α (**a3**), β (**a4**), γ_L_ (**a5**), γ_H_ (**a6**). For a complete description of PGI areas, see also Supplementary Table S8-S13 as some subcortical regions are not depicted.

In sum, we found PGI-assortative nodes in frontal areas for α and β rhythms, and in posterior areas, mainly occipital and temporal, for both γ oscillations. PGI-disassortative nodes was predominant in the cingulate cortex in β and γ_H_ (also PCC in β band), precuneus and some parietal areas (β). Finally, slow rhythms were characterized by PGI-core-peripheriness mostly in the occipital, temporal (δ) and frontal regions (θ), as well as small portion of the frontal lobe (β).

## 4 Discussion

To date, the features of human brain meso-scale structure during resting state have not been fully investigated. Specifically, meso-scale spectral features are still largely unexplored, and evidence about how the repertoire of community motif (i.e. assortative, disassortative and core-periphery) is associated with different neural oscillations is missing. Thus, we aimed at filling this knowledge gap by using WSBM to infer the diversity of the latent community structure, estimated from source-reconstructed hdEEG resting state signals. By relying on the spectral richness of hdEEG recordings, we could investigate how FC network meso-scale is organized across frequency bands. We described the spatial distribution of communities and we characterized their interactions across the frequency domain. According to our results, the meso-scale is characterized by a frequency-specific organization. Also, with our experiments and analyses, we highlighted a relevant amount of non-assortative community structure. Additionally, we found that specific brain areas preferably participate in specific community motifs, depending on the oscillatory rhythms. Finally, we identified the so-called PGI areas exhibiting invariance across partitions and the highest community motif level.

### 4.1 Meso-scale structure is frequency-dependent

We observed an increasing occurrence of the assortative structure when increasing the neuronal oscillation frequency from δ to γ_H_, with greater values in the γ_L_ band. Conversely, the disassortative structure showed an opposite trend, exhibiting highest values in δ and θ slow rhythms. Importantly, communities mostly followed a core-periphery structure to interact among each other, regardless of the specific frequency band. Indeed, core-periphery was uniformly distributed across the spectrum, without any particular trend. Given this frequency-specific organization, we propose in the following that each of the three meso-scale structure might underlie a particular mechanism of neuronal oscillations.

Slow brain rhythms, are characterized by long-range communication (Leong et al., 2016), and information is exchanged over long-distance. We suggest that this behavior might be expressed by the disassortative structure which is higher in δ and θ when compared to the γ_L_ rhythm, thus favoring integration between long-distance areas belonging to spatially distinct modules (Betzel, Bertolero, et al., 2018). Therefore, we can consider the disassortative structure as a meso-scale property of the slow oscillations. Moreover, in δ and θ bands, the WSBM clustering is more fragmented than in the higher frequency bands, likely because these oscillatory regimens are characterized by long-range interactions (Leong et al., 2016), which might complicate the formation of communities whose brain nodes are spatially closed and/or belong to the same functional domain. This communities’ shattering in slow rhythms can be observed at each of the K-th partition we performed (see Figure 1 and Figure S1-S6) and might further support the hypothesis of central role of disassortativity for slow oscillations.

On the other hand, for higher frequencies (from δ to γ_L_ and γ_H_ band) we encountered not only an increase of the assortative meso-scale structure (with a significant peak in γ_L_), but also a clearer subdivision of the communities with respect to the low bands (see Figure 1) that, in turn, may reflect a local and spatially– segregated processing. In further support of this, there are few branches deviating from the originating community for the high rhythms (see Figure 4), further corroborating the idea that, for fast oscillations, the community detection found more segregated clusters. In fact, gamma oscillations might represent a rhythmic synaptic inhibition mediated by parvalbumin-expressing inhibitory interneurons and the interconnected pyramidal neurons (Buzsaki, Logothetis, & Singer, 2013; Sohal, 2012, 2016). Gamma-oscillations and the associated assortative structure, might thus resemble a local processing of areas that are spatially contiguous and/or share the same function during resting state.

Core-periphery community structure, which is the most represented in all the frequency bands, may underlie the meso-scale backbone supporting two well-known strictly interconnected principles of brain organization: local segregation and functional integration (Sporns, 2010). The highly dense core could represent the segregation while the interactions between the core and the nodes located in the peripheries could indicate the presence of functional connections, which might in turn reflect an efficient integration. From this perspective, a plausible interpretation is that the core-periphery meso-scale structure might be a good candidate to support this physiological balance between segregation and integration (Sporns, 2010) across frequency bands. Previous studies have indeed demonstrated that neuronal oscillations in the gamma band reflect not only a local processing, but may also exhibit synchronization across long-distance areas (Buzsáki & Schomburg, 2015; Sohal, 2016).

However, all these interpretations have a speculative nature and further studies, that may infer meso-scale structure from effective connectivity (EC) or from combined information of FC/EC with anatomical connectivity from MRI (Glomb et al., 2019), are needed to potentially validate these claims.

In summary, we found the emergence of non-assortative meso-scale structures. According to previous studies focusing on MRI and fMRI, the brain presents a mixed meso-scale organization, but the network predominantly exhibits modular/assortative meso-scale structures, specifically during resting state (Betzel, Bertolero, et al., 2018; Betzel, Medaglia, et al., 2018) and, to a lesser extent, during cognitive tasks (Betzel, Bertolero, et al., 2018). Our analysis showed that the amount of assortative modules was reduced when decomposing the time-course in the frequency domain, and a clear non-assortative organization arose.

### 4.2 Community motifs rely on brain areas location and frequency band

We further aimed at identifying which brain regions were at the same time insensitive to partitioning and exhibited the highest community motifs in each band (PGI areas). PGI-assortative areas were located, for γ oscillations, in posterior areas spanning portions of the occipital and temporal lobes, where they potentially resembled the local processing that is peculiar of γ oscillations (see above). Furthermore, some of these posterior areas, were the visual cortices and the inferior temporal lobe, which are components of the visual what/ventral stream, associated with object recognition (Reddy & Kanwisher, 2006). Interestingly, our subjects were fixating a cross in the middle of a computer screen (eyes-open resting state paradigm). By contrast, some of the PGI-disassortative areas were located in PCC and angular gyrus (in β band), which are well-known areas of the default mode network (DMN) (Raichle, 2015). In particular, it has been demonstrated that in β and γ rhythms there are specific connectivity profile between the areas of the DMN (Samogin et al., 2019), including PCC and angular gyrus. Again in the β band, the precuneus showed high disassortativity and this area has been usually associated to self-referential processing (Cavanna & Trimble, 2006), but it has been also identified as another key area of the DMN, according to fMRI study (Utevsky, Smith, & Huettel, 2014). In sum, these DMN areas are part of communities interacting disassortatively, and this might underlie their integration nature (Betzel, Bertolero, et al., 2018).

Frontal areas in α and β showed significant PGI-assortativity. Some of these frontal areas are part of the dorsal-lateral prefrontal cortex that is a pivotal area involved in several high-order executive functions (Barbey, Colom, & Grafman, 2013), such as decision making (Heekeren, Marrett, Ruff, Bandettini, & Ungerleider, 2006) and working memory (Lara & Wallis, 2015), among others (Miller & Cohen, 2001).

Regions showing significant core-peripheriness are instead typical of slow oscillations in frontal lobe (θ) and in occipital and temporal lobes (δ). Particularly, these areas were assortative in higher bands: posterior areas in gamma bands and frontal areas in β. We therefore posit that the same regions might exhibit a community motif depending on the specific neuronal oscillation.

### 4.3 Study limitations

One possible limitation of our study is that we based our analysis on measurements of spontaneous brain activity, which is brain activity generated in the absence of an explicit task. However, we must recall that many studies about resting state paradigm supported the interpretation that plasticity mechanisms, which depend on previous subjects’ interaction with the external world (Carrillo-Reid, Miller, Hamm, Jackson, & Yuste, 2015; Guerra-Carrillo, Mackey, & Bunge, 2014; Northoff, 2016), sculpt the activity and bridge together neurons belonging to the same functional domain (motor, visual, auditory, default mode network, among others) (Kelly & Castellanos, 2014; Vincent et al., 2007). Hence, also during spontaneous activity, the co-activation of these neurons is facilitated and successfully detected by resting state processing pipelines, independently from the non-invasive neuroimaging dataset (Coquelet et al., 2020; Liu et al., 2017). Thus, upon this framework, albeit no task is being performed, we can say that the spontaneous activity organization at the meso-scale level, might largely reflect experience-dependent plasticity mechanisms, posing the above-described interpretation as possible candidate of how the brain shapes information flow.

An additional observation refers to the number of communities K, which is a free parameter in the context of community detection by WSBM. Here, it has been varied across a range of partitions, allowing for multiple-grains meso-scale analysis. This examination showed that each of the frequency might exhibit its own optimal number of communities. In support of this claim, the flows’ reconfiguration in Figure 4 indicates that each band has its own reconfiguration pattern over the partitions, possibly suggesting, in particular for the lower bands, a sub-optimal fragmentation of modules (when the partition number increases). Moreover, modules fade away and/or remained invariant, depending on the frequency band of interest. Therefore, we suggest that future works should also focus on finding the best K per each of the band considered, by performing appropriate parameter selection procedures.

## 5 Conclusion

Our analysis allowed to observe WSBM-estimated meso-scale organization with a different focus: by investigating FC in different frequency bands, we captured peculiar features of module interactions revealing the non-assortative nature of resting state networks, demonstrating its frequency-specificity. Furthermore, this study showed that WSBM applied to sources-level neuronal oscillations is able to reveal yet unknown properties of FC topological organization.

Overall, these results may be taken into consideration for future studies that will address the pathophysiological mechanisms underlying neurological/psychiatric disorders (Babiloni et al., 2019; Siegel et al., 2012). It would indeed be crucial to examine how the presence of a neurological disease can affect the meso-scale structure and whether and how a neurorehabilitation intervention can impact the re-organization of brain networks and the interactions among communities. This will have a direct impact in the clinical assessment of sensory, motor and cognitive functions, being EEG acquisitions widely employed in the clinical setting. Collectively, the results of our study advance the network neuroscience field by highlighting new features of brain meso-scale organization.

## Acknowledgments

The authors would like to thank the Rehab Technologies IIT-INAIL joint lab led by Dr. Lorenzo De Michieli for supporting the conducted research. The authors would like to thank Dr. Gaia Bonassi, Dr. Mingqi Zhao and Federico Barban for collecting part of the experimental data used in this work and to Dr. Stefano Buccelli for discussion on methodology. The authors are grateful to Silvia Chiappalone for assistance with the graphics and Samuel Stedman for proofreading the manuscript.

## Author contribution

R.I., M.C. conceived the study. R.I., M.S. collected the data. R.I., M.S, D.S., S.B. and M.C. designed the methods. R.I., M.S. performed the analysis. R.I., M.S., D.M., L.A. and M.C. interpreted and discussed the results. R.I., M.S., M.C. prepared the figures. R.I. wrote the first version of the manuscript. All the authors contributed to the revision of the manuscript. Part of this work was supported by the Gossweiler foundation, granted to L. Avanzino (PI), D. Mantini, and M. Chiappalone.

## Data availability statement

We plan to make the EEG data sets available on Mendeley-Data repository (or similar). Currently, data are being used for another publication of co-authors of ours and we cannot share the EEG data at this point.

## Supplementary Materials

### 5.1 T1-weighted structural images acquisition

Subjects underwent T1-weighted using either a 3T or 1.5 T scanners. See Supplementary Table 1 for the T1-weighted acquisition parameters.

**Table S1.**

| Number of subject | 19 | 6 | 4 |
| --- | --- | --- | --- |
| MRI scanner | 3T Achieva<br>(Philips Medical System,<br>The Netherlands) | 3 T MAGNETOM Prisma<br>(Siemens AG, Healthcare<br>Sector, Germany) | 1.5 T Signa Excite<br>(General Electric<br>Healthcare, USA) |
| T1-weighted sequence name | Magnetization Prepared Rapid Acquisition Gradient Echo (MPRAGE) | Magnetization Prepared Rapid Acquisition Gradient Echo (MPRAGE) | Fast Spoiled Gradient Echo (SPGR) |
| Coil | 32-channel head coil | 32-channel head coil | 8-channels phased array head coil |
| Voxel Size | 0.98×0.98×1.2 mm<sup>3</sup> | 1×1×1 mm<sup>3</sup> | 1×1×1 mm<sup>3</sup> |
| TR/TE | 9.6/4.6 ms | 2.3/2.96 ms | 11.8/5.18 ms |
| Field of View (F.O.V) | 250x250 mm<sup>2</sup> | 256×256 mm<sup>2</sup> | 256×256 mm<sup>2</sup> |

### 5.2 Meso-scale structure K = 3

**Fig. S1.**
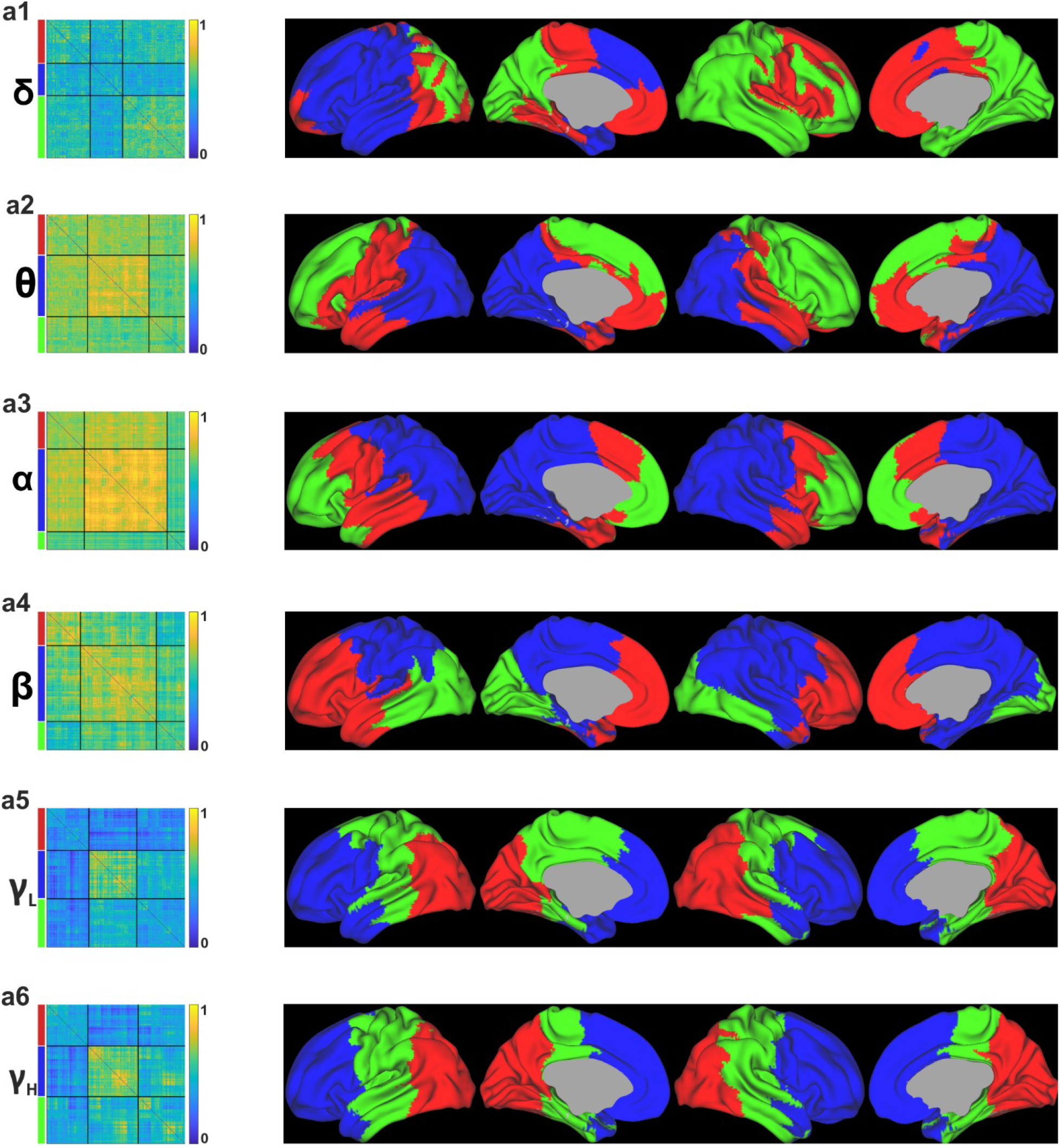
Organization of the meso-scale structure in the frequency domain. Each row represents the K = 3 community assignments in each of the considered frequency band: δ (**a1**), θ (**a2**), α (**a3**), β (**a4**), γ_L_ (**a5**), γ_H_ (**a6**). Each row contains the re-ordered group level adjacency matrix after WSBM estimation and spatial distribution of the three partitions across the brain. Colors on left side of each adjacency matrix match with the colors overlaid on the brain.

### 5.3 Meso-scale structure K = 4

**Fig. S2.**
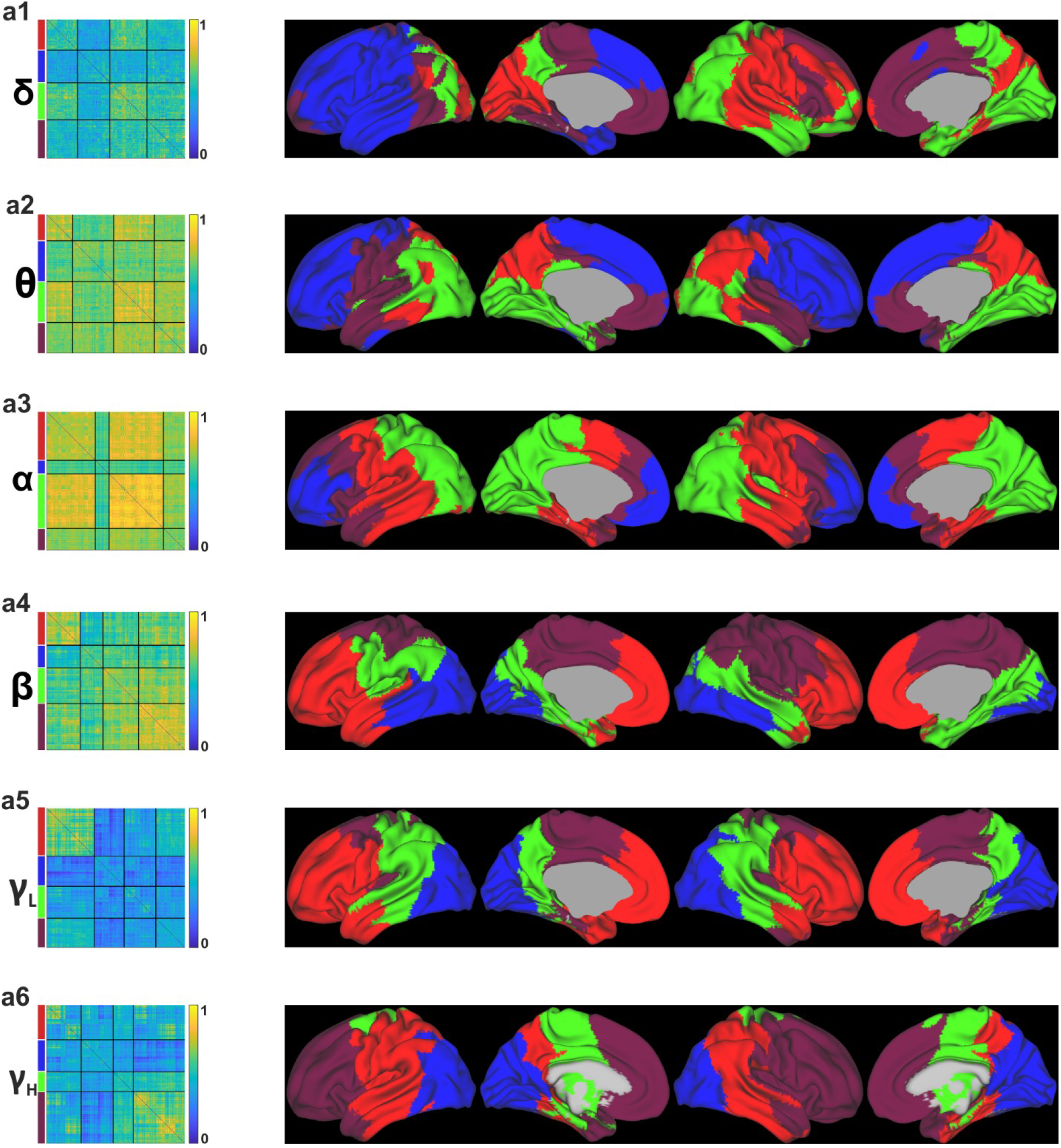
Organization of the meso-scale structure in the frequency domain. Each row represents the K = 4 community assignments in each of the considered frequency band: δ (**a1**), θ (**a2**), α (**a3**), β (**a4**), γ_L_ (**a5**), γ_H_ (**a6**). Each row contains the re-ordered group level adjacency matrix after WSBM estimation and spatial distribution of the four partitions across the brain. Colors on left side of each adjacency matrix match with the colors overlaid on the brain.

### 5.4 Meso-scale structure K = 6

**Fig. S3.**
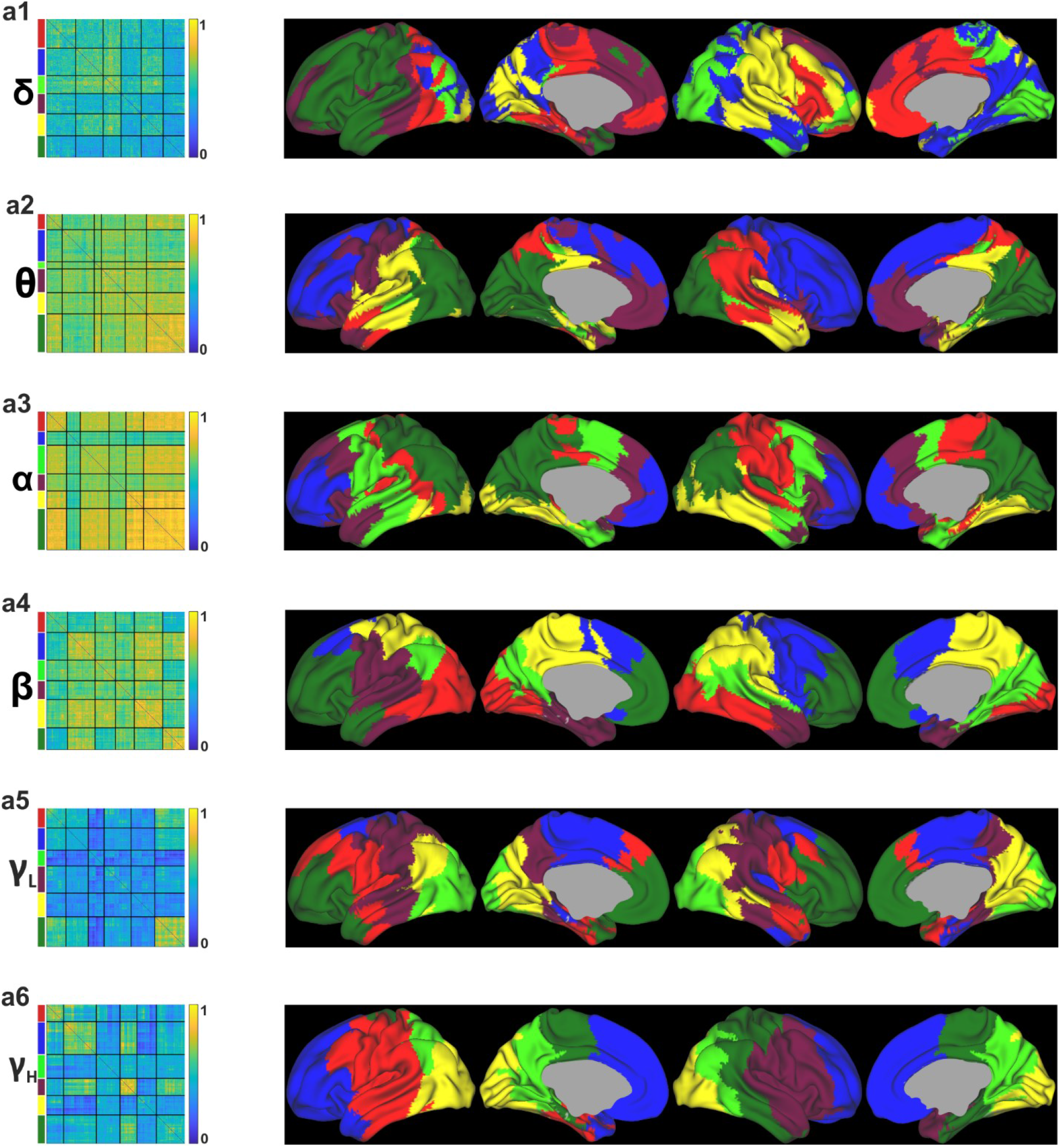
Organization of the meso-scale structure in the frequency domain. Each row represents the K = 6 community assignments in each of the considered frequency band: δ (**a1**), θ (**a2**), α (**a3**), β (**a4**), γ_L_ (**a5**), γ_H_ (**a6**). Each row contains the re-ordered group level adjacency matrix after WSBM estimation and spatial distribution of the six partitions across the brain. Colors on left side of each adjacency matrix match with the colors overlaid on the brain.

### 5.5 Meso-scale structure K = 7

**Fig. S4.**
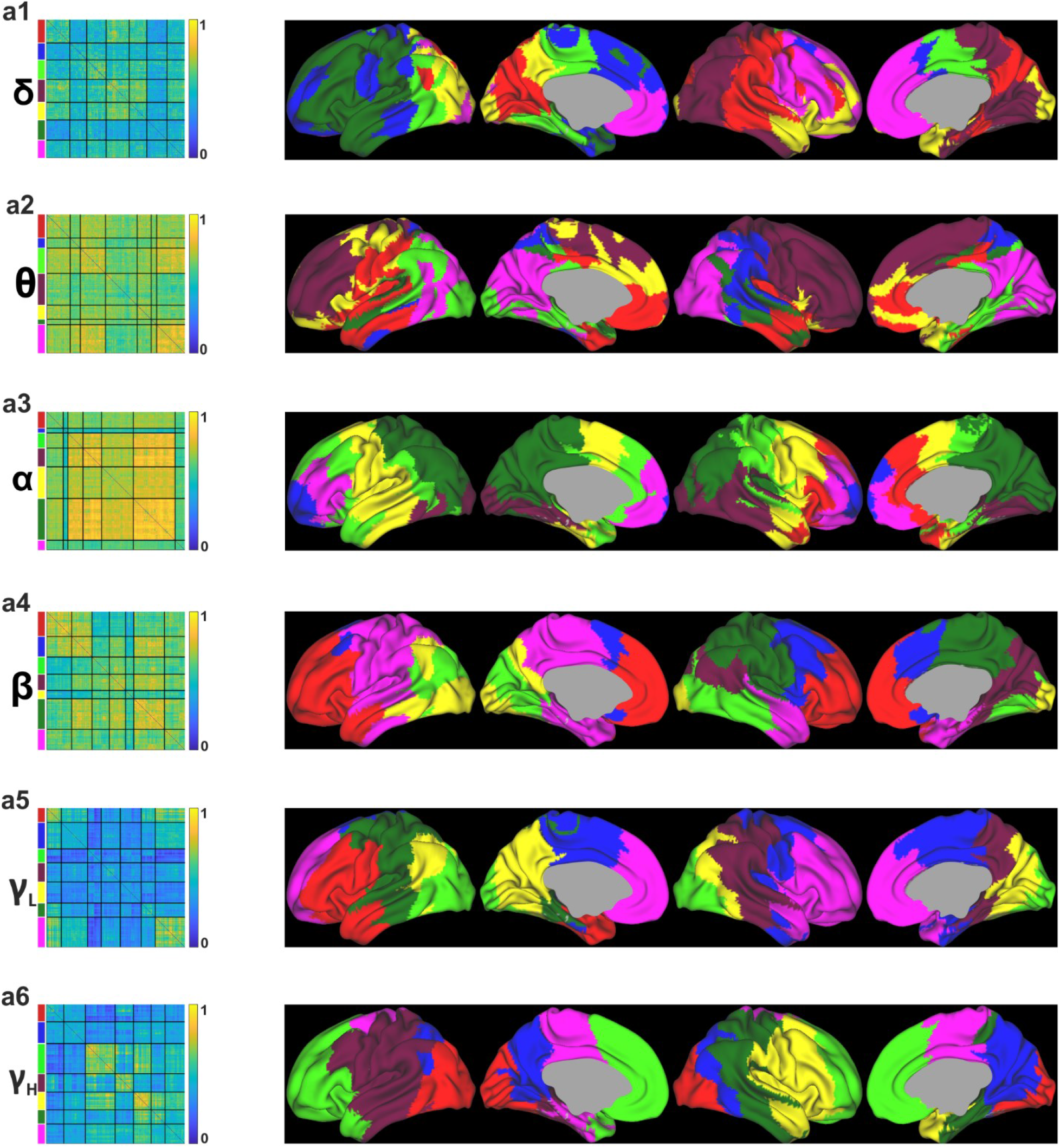
Organization of the meso-scale structure in the frequency domain. Each row represents the K = 7 community assignments in each of the considered frequency band: δ (**a1**), θ (**a2**), α (**a3**), β (**a4**), γ_L_ (**a5**), γ_H_ (**a6**). Each row contains the re-ordered group level adjacency matrix after WSBM estimation and spatial distribution of the seven partitions across the brain. Colors on left side of each adjacency matrix match with the colors overlaid on the brain.

### 5.6 Meso-scale structure K = 8

**Fig. S5.**
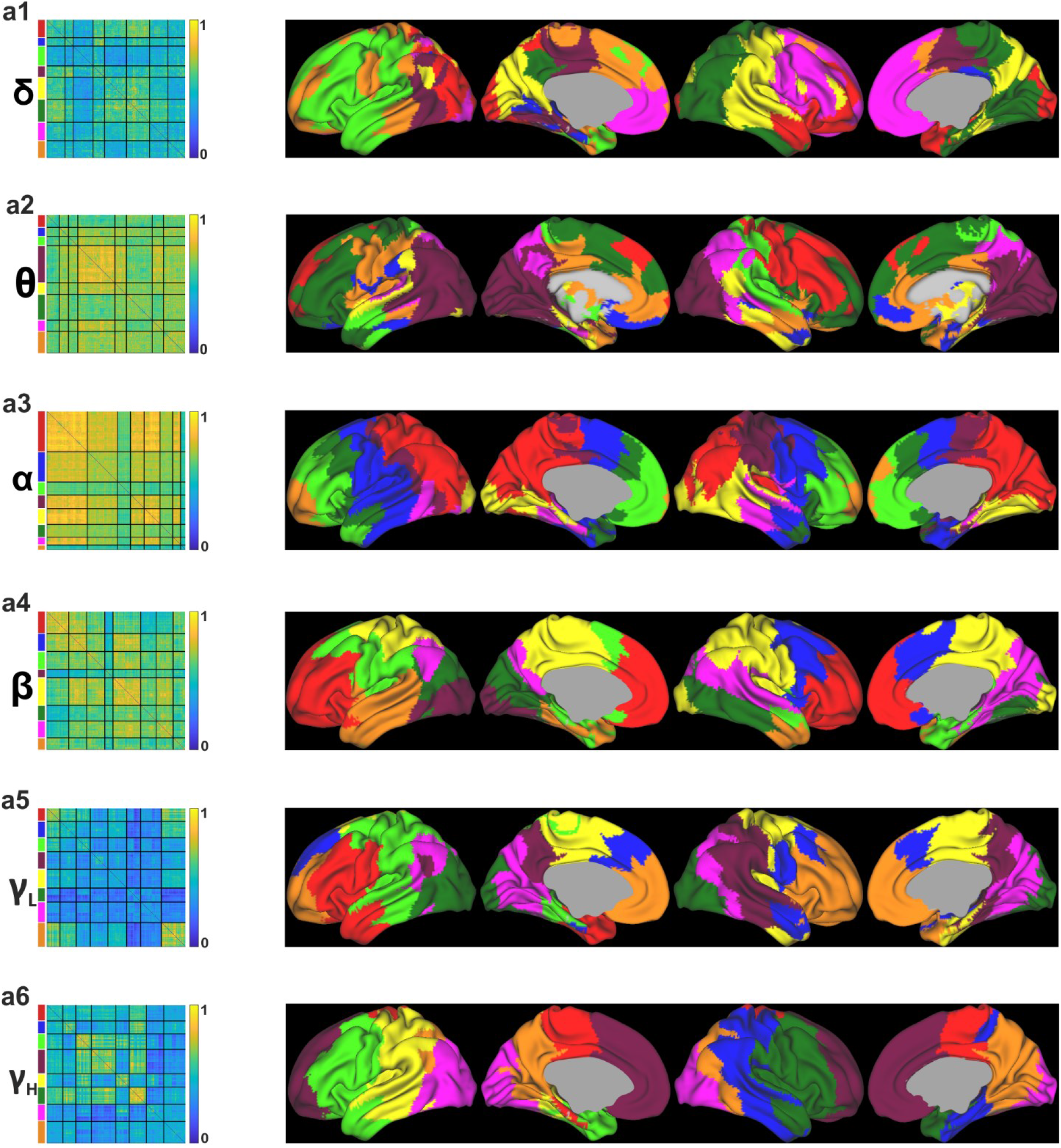
Organization of the meso-scale structure in the frequency domain. Each row represents the K = 8 community assignments in each of the considered frequency band: δ (**a1**), θ (**a2**), α (**a3**), β (**a4**), γ_L_ (**a5**), γ_H_ (**a6**). Each row contains the re-ordered group level adjacency matrix after WSBM estimation and spatial distribution of the eight partitions across the brain. Colors on left side of each adjacency matrix match with the colors overlaid on the brain.

### 5.7 Meso-scale interactions per each K-value

**Fig. S6.**
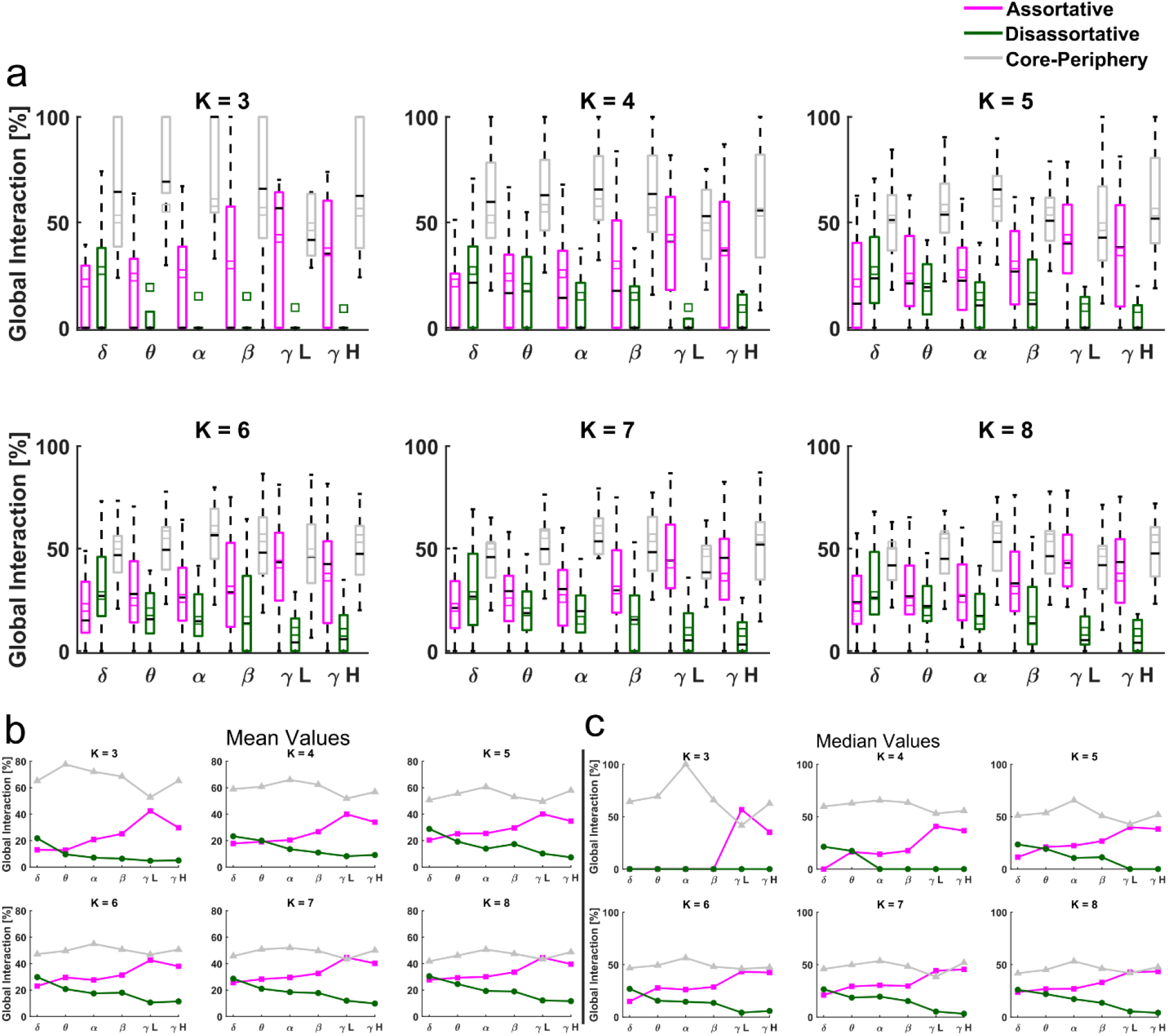
Frequency-specificity of meso-scale-structure at each of the K partition considered. a. Each subpanels show boxplots (at a fixed partition K) with distributions across participants of the three meso-scale motifs per each frequency band. Boxplots show upper and lower bound of the distributions at 25^th^ and 75^th^ percentile. Whiskers extend to the most extreme data points not considered outliers. The black horizontal lines represent the median, while the small colored squares indicate the mean of the distributions. **b-c**. Mean (b) and median (c) values of each meso-scale structure distributions across frequency bands.

### 5.8 Significant areas after non parametric testing 827

The following tables show, frequency by frequency, the significant node’s name, MNI coordinate, hemisphere (according to the AICHA atlas), and global community motifs values (assortative or disassortative or core-periphery). Part of the significant nodes that are reported are the same which are overlaid onto the T1-weighted template in Figure 3 panel b of the main manuscript. Other nodes are here represented, but not in the Figure 3, as they belong to subcortical areas. Colors resembles those of the Figure 2 (pink: assortative, green: disassortative and gray: core-periphery).

**Table S2.** List of significant regions for δ band (p < 0.01). Color code are Pink: Assortative; Green: Disassortative; Gray: Core-periphery.

| region | x | y | z | side | value [%] |
| --- | --- | --- | --- | --- | --- |
| S Sup Temporal 5 L | 0 | -58 | 26 | L | 26.122 |
| G Precuneus 3 R | 6 | -52 | 32 | R | 26.232 |
| G Precuneus 4 L | -7 | -49 | 45 | L | 26.523 |
| G Precuneus 8 L | -7 | -71 | 40 | L | 25.877 |
| N Caudate 5 L | -13 | 10 | 8 | L | 26.216 |
| S Sup Frontal 2 R | 28 | 56 | 7 | R | 33.602 |
| G Insula anterior 1 R | 19 | 7 | -19 | R | 33.363 |
| G Insula anterior 2 R | 35 | 18 | -13 | R | 32.911 |
| G Insula anterior 4 R | 41 | 15 | 4 | R | 32.563 |
| G Insula anterior 5 R | 42 | 2 | -9 | R | 35.398 |
| S Sup Temporal 1 R | 52 | 13 | -26 | R | 33.514 |
| S Sup Temporal 2 R | 54 | -2 | -15 | R | 32.932 |
| G Temporal Pole Sup 1 R | 36 | 16 | -24 | R | 33.183 |
| G Temporal Pole Sup 2 R | 44 | 9 | -16 | R | 34.974 |
| G Temporal Pole Mid 1 R | 48 | 8 | -33 | R | 33.199 |
| G Hippocampus 1 R | 30 | -5 | -18 | R | 33.61 |
| G ParaHippocampal 1 R | 14 | -4 | -18 | R | 32.794 |
| N Amygdala 1 R | 21 | 2 | -12 | R | 34.42 |
| N Caudate 6 R | 15 | 7 | 18 | R | 34.666 |
| N Putamen 2 R | 23 | 7 | 1 | R | 35.099 |
| N Putamen 3 R | 29 | -4 | 1 | R | 35.995 |
| N Thalamus 1 L | -4 | 0 | 1 | L | 33.555 |
| N Thalamus 1 R | 4 | 0 | 1 | R | 33.736 |
| N Thalamus 3 L | -3 | -7 | -1 | L | 33.495 |
| N Thalamus 3 R | 3 | -7 | -1 | R | 32.622 |
| N Thalamus 4 L | -3 | -14 | 8 | L | 32.514 |
| N Thalamus 4 R | 3 | -14 | 9 | R | 32.976 |
| N Thalamus 9 L | -5 | -11 | -7 | L | 33.681 |
| N Thalamus 9 R | 5 | -10 | -6 | R | 32.745 |
| G Frontal Mid 5 R | 42 | 17 | 41 | R | 58.778 |
| S Precentral 4 R | 44 | 1 | 48 | R | 59.154 |
| S Precentral 5 R | 57 | 8 | 29 | R | 59.501 |
| G Angular 2 L | -38 | -70 | 39 | L | 58.469 |
| G Occipital Pole 1 L | -21 | -96 | -14 | L | 60.123 |
| G Occipital Pole 1 R | 24 | -93 | -13 | R | 59.554 |
| G Occipital Lat 1 R | 17 | -94 | 4 | R | 59.596 |
| G Occipital Lat 2 L | -26 | -94 | -1 | L | 58.788 |
| G Occipital Lat 2 R | 28 | -89 | -2 | R | 63.474 |
| G Occipital Lat 2 L | -40 | -84 | -12 | L | 59.39 |
| G Occipital Lat 2 R | 43 | -81 | -10 | R | 61.601 |
| G Occipital Lat 4 L | -31 | -89 | 8 | L | 60.298 |
| G Occipital Lat 4 R | 34 | -85 | 9 | R | 58.989 |
| G Occipital Lat 5 R | 36 | -76 | 2 | R | 60.665 |
| G Occipital Mid 3 R | 45 | -63 | 15 | R | 59.491 |
| G Occipital Inf 1 L | -48 | -69 | -4 | L | 58.864 |
| G Occipital Inf 1 R | 50 | -60 | -9 | R | 58.298 |
| G Temporal Sup 1 L | -55 | -1 | 2 | L | 58.762 |
| S Sup Temporal 5 R | 51 | -54 | 21 | R | 58.786 |
| G Temporal Mid 1 L | -61 | -12 | -17 | L | 58.125 |
| G Temporal Mid 4 L | -53 | -59 | 7 | L | 58.585 |
| G Temporal Mid 4 R | 57 | -53 | 3 | R | 58.7 |
| G Temporal Inf 1 L | -47 | -15 | -33 | L | 58.406 |
| G Temporal Inf 2 L | -59 | -26 | -24 | L | 58.756 |
| G Temporal Inf 4 R | 54 | -58 | -11 | R | 58.373 |
| G Temporal Inf 5 L | -45 | -64 | 6 | L | 60.159 |
| G Lingual 5 R | 11 | -79 | -7 | R | 58.69 |
| G Fusiform 7 R | 23 | -80 | -8 | R | 60.951 |

**Table S3.** List of significant regions for θ band (p < 0.01). Color code are Pink: Assortative; Green: Disassortative; Gray: Core-periphery.

| region | x | y | z | side | value [%] |
| --- | --- | --- | --- | --- | --- |
| G Occipital Lat 2 L | -40 | -84 | -12 | L | 24.28 |
| G ParaHippocampal 1 R | 14 | -4 | -18 | R | 24.117 |
| N Thalamus 3 L | -3 | -7 | -1 | L | 23.622 |
| N Thalamus 4 L | -3 | -14 | 8 | L | 24.332 |
| N Thalamus 4 R | 3 | -14 | 9 | R | 23.992 |
| N Thalamus 5 R | 13 | -17 | 6 | R | 24.858 |
| N Thalamus 7 R | 9 | -26 | 10 | R | 23.949 |
| N Thalamus 8 R | 4 | -25 | 13 | R | 24.392 |
| G Frontal Sup 2 L | -12 | 47 | 40 | L | 63.385 |
| S Sup Frontal 1 R | 20 | 63 | -6 | R | 62.651 |
| S Sup Frontal 2 L | -27 | 56 | 1 | L | 62.47 |
| G Frontal Mid 1 L | -40 | 41 | 20 | L | 64.127 |
| G Frontal Mid 2 L | -33 | 44 | 25 | L | 62.507 |
| G Frontal Mid 4 R | 39 | 22 | 46 | R | 63.049 |
| S Inf Frontal 1 L | -44 | 38 | 12 | L | 65.142 |
| G Frontal Inf Tri 1 L | -49 | 26 | 5 | L | 62.997 |
| G Frontal Mid Orb 1 L | -35 | 57 | -8 | L | 64.087 |
| G Frontal Mid Orb 2 L | -41 | 49 | -5 | L | 63.498 |
| S Precentral 2 R | 26 | -8 | 58 | R | 64.44 |
| S Precentral 2 R | 20 | -6 | 68 | R | 63.224 |
| G Insula anterior 2 L | -34 | 24 | 1 | L | 63.498 |
| G Insula anterior 4 L | -41 | 15 | 3 | L | 62.574 |
| G Supp Motor Area 2 R | 10 | 19 | 61 | R | 62.946 |

**Table S4.** List of significant regions for α band (p < 0.01). Color code are Pink: Assortative; Green: Disassortative; Gray: Core-periphery.

| region | x | y | z | side | value [%] |
| --- | --- | --- | --- | --- | --- |
| G Frontal Sup 1 L | -16 | 65 | 13 | L | 32.407 |
| G Frontal Sup 1 R | 13 | 68 | 11 | R | 32.579 |
| G Frontal Sup 2 L | -12 | 47 | 40 | L | 31.16 |
| S Sup Frontal 1 L | -22 | 61 | -8 | L | 32.611 |
| S Sup Frontal 1 R | 20 | 63 | -6 | R | 31.672 |
| S Sup Frontal 2 L | -27 | 56 | 1 | L | 32.708 |
| G Frontal Mid 1 L | -40 | 41 | 20 | L | 31.779 |
| S Inf Frontal 1 L | -44 | 38 | 12 | L | 31.752 |
| G Frontal Mid Orb 1 L | -35 | 57 | -8 | L | 32.727 |
| G Frontal Mid Orb 1 R | 36 | 57 | -6 | R | 31.836 |
| G Frontal Mid Orb 2 L | -41 | 49 | -5 | L | 32.467 |
| G Frontal Inf Orb 1 L | -42 | 31 | -17 | L | 33.187 |
| S Orbital 2 L | -31 | 34 | -13 | L | 31.78 |
| G Frontal Sup Medial 1 L | -8 | 52 | 12 | L | 31.46 |
| G Frontal Sup Medial 2 L | -7 | 53 | 25 | L | 32.12 |
| G Frontal Sup Medial 2 R | 7 | 57 | 25 | R | 31.66 |
| S Anterior Rostral 1 L | -7 | 47 | -3 | L | 32.204 |
| G Frontal Med Orb 1 L | -10 | 65 | -2 | L | 31.897 |
| G Frontal Med Orb 1 R | 7 | 65 | -3 | R | 32.411 |
| G Frontal Med Orb 2 L | -5 | 49 | -12 | L | 32.084 |
| G Cingulum Ant 1 L | -6 | 37 | 3 | L | 31.46 |
| G ParaHippocampal 3 R | 18 | -14 | -29 | R | 19.858 |
| G Fusiform 1 R | 31 | -8 | -34 | R | 20.471 |
| N Amygdala 1 R | 21 | 2 | -12 | R | 19.148 |
| N Caudate 6 L | -16 | 4 | 19 | L | 19.658 |
| N Caudate 7 L | -18 | -12 | 25 | L | 19.304 |
| N Thalamus 2 L | -9 | -9 | 13 | L | 19.154 |

**Table S5.** List of significant regions for β band (p < 0.01). Color code are Pink: Assortative; Green: Disassortative; Gray: Core-periphery.

| region | x | y | z | side | value [%] |
| --- | --- | --- | --- | --- | --- |
| G Frontal Sup 2 L | -12 | 47 | 40 | L | 35.222 |
| S Sup Frontal 1 L | -22 | 61 | -8 | L | 36.377 |
| S Sup Frontal 2 L | -27 | 56 | 1 | L | 35.639 |
| S Sup Frontal 2 L | -26 | 43 | 29 | L | 35.617 |
| G Frontal Mid 1 L | -40 | 41 | 20 | L | 35.427 |
| G Frontal Mid Orb 1 L | -35 | 57 | -8 | L | 36.193 |
| G Frontal Mid Orb 2 L | -41 | 49 | -5 | L | 36.738 |
| G Parietal Sup 2 L | -30 | -51 | 64 | L | 19.628 |
| G Angular 2 R | 45 | -62 | 36 | R | 19.62 |
| S Intraparietal 2 R | 37 | -52 | 48 | R | 20.495 |
| S Sup Temporal 5 L | 0 | -58 | 26 | L | 20.743 |
| S Cingulate 6 R | 11 | -34 | 46 | R | 20.016 |
| S Cingulate 7 R | 9 | -41 | 61 | R | 19.999 |
| G Cingulum Post 2 L | -4 | -39 | 27 | L | 19.57 |
| G Cingulum Post 2 R | 8 | -43 | 31 | R | 19.747 |
| G Precuneus 1 R | 13 | -53 | 14 | R | 19.681 |
| G Precuneus 2 R | 5 | -56 | 20 | R | 20.661 |
| G Precuneus 3 L | -7 | -51 | 34 | L | 20.463 |
| G Precuneus 3 R | 6 | -52 | 32 | R | 21.635 |
| G Precuneus 4 L | -7 | -49 | 45 | L | 20.962 |
| G Precuneus 4 R | 8 | -48 | 43 | R | 21.019 |
| G Precuneus 5 L | -8 | -55 | 55 | L | 20.64 |
| G Precuneus 5 R | 7 | -50 | 53 | R | 20.467 |
| G Precuneus 7 R | 7 | -63 | 36 | R | 21.24 |
| G Precuneus 8 R | 11 | -68 | 41 | R | 20.624 |
| S Parietooccipital 2 R | 16 | -61 | 26 | R | 20.308 |
| S Parietooccipital 3 R | 14 | -73 | 37 | R | 20.788 |
| G Calcarine 1 R | 12 | -69 | 10 | R | 19.943 |
| N Thalamus 7 R | 9 | -26 | 10 | R | 19.674 |
| G Frontal Inf Tri 1 R | 50 | 29 | 5 | R | 61.479 |
| G Frontal Mid Orb 2 R | 40 | 50 | -4 | R | 62.152 |
| G Temporal Pole Mid 2 R | 35 | 12 | -34 | R | 61.546 |
| G Temporal Pole Mid 3 L | -26 | 6 | -37 | L | 61.284 |

**Table S6.** List of significant regions for γ_L_ band (p < 0.01). Color code are Pink: Assortative; Green: Disassortative; Gray: Core-periphery.

| region | x | y | z | side | value [%] |
| --- | --- | --- | --- | --- | --- |
| G Angular 2 R | 47 | -63 | 25 | R | 53.027 |
| S Intraoccipital 1 R | 28 | -69 | 33 | R | 51.282 |
| G Occipital Pole 1 R | 24 | -93 | -13 | R | 54.388 |
| G Occipital Lat 1 L | -15 | -97 | 1 | L | 50.661 |
| G Occipital Lat 1 R | 17 | -94 | 4 | R | 53.358 |
| G Occipital Lat 2 R | 28 | -89 | -2 | R | 54.47 |
| G Occipital Lat 2 R | 43 | -81 | -10 | R | 53.198 |
| G Occipital Lat 4 R | 34 | -85 | 9 | R | 52.965 |
| G Occipital Lat 5 R | 36 | -76 | 2 | R | 54.258 |
| G Occipital Sup 1 R | 18 | -81 | 37 | R | 53.385 |
| G Occipital Sup 2 R | 17 | -88 | 26 | R | 56.233 |
| G Occipital Mid 1 R | 36 | -74 | 25 | R | 55.186 |
| G Occipital Mid 2 R | 41 | -73 | 12 | R | 53.978 |
| G Occipital Mid 3 R | 45 | -63 | 15 | R | 54.502 |
| G Occipital Mid 4 R | 41 | -74 | 30 | R | 51.026 |
| G Occipital Inf 1 L | -48 | -69 | -4 | L | 50.285 |
| G Occipital Inf 1 R | 50 | -60 | -9 | R | 58.452 |
| G Occipital Inf 2 L | -45 | -71 | -7 | L | 50.025 |
| G Occipital Inf 2 R | 47 | -65 | -7 | R | 58.103 |
| S Sup Temporal 3 R | 53 | -32 | 0 | R | 50.679 |
| G Temporal Mid 3 R | 62 | -31 | -5 | R | 49.579 |
| G Temporal Mid 4 R | 57 | -53 | 3 | R | 54.639 |
| G Temporal Inf 3 R | 57 | -46 | -14 | R | 56.333 |
| G Temporal Inf 4 L | -50 | -61 | -8 | L | 50.74 |
| G Temporal Inf 4 R | 54 | -58 | -11 | R | 59.635 |
| G Temporal Inf 5 R | 49 | -58 | 4 | R | 56.76 |
| S Parietooccipital 4 R | 21 | -59 | 6 | R | 52.713 |
| S Parietooccipital 5 R | 21 | -66 | 20 | R | 51.374 |
| S Parietooccipital 6 R | 15 | -73 | 29 | R | 50.283 |
| G Cuneus 1 R | 9 | -80 | 29 | R | 51.556 |
| G Cuneus 2 L | -4 | -92 | 19 | L | 50.12 |
| G Cuneus 2 R | 8 | -89 | 19 | R | 53.465 |
| G Calcarine 1 R | 12 | -69 | 10 | R | 50.712 |
| G Calcarine 2 R | 10 | -78 | 9 | R | 51.219 |
| G Calcarine 3 L | -7 | -95 | -1 | L | 50.133 |
| G Calcarine 3 R | 11 | -94 | 1 | R | 52.221 |
| G Lingual 2 R | 21 | -60 | -6 | R | 51.687 |
| G Lingual 3 R | 13 | -64 | -2 | R | 49.91 |
| G Lingual 4 R | 13 | -72 | -9 | R | 52.965 |
| G Lingual 5 R | 11 | -79 | -7 | R | 53.077 |
| G Lingual 6 R | 7 | -79 | -3 | R | 52.147 |
| G Fusiform 2 R | 38 | -25 | -24 | R | 49.763 |
| G Fusiform 3 R | 37 | -31 | -24 | R | 51.545 |
| G Fusiform 4 R | 44 | -46 | -18 | R | 54.387 |
| G Fusiform 5 R | 32 | -47 | -11 | R | 51.589 |
| G Fusiform 6 R | 29 | -62 | -9 | R | 54.249 |
| G Fusiform 7 R | 23 | -80 | -8 | R | 53.935 |
| G Parietal Sup 2 L | -30 | -51 | 64 | L | 12.722 |
| G Parietal Sup 5 L | -16 | -61 | 59 | L | 12.577 |
| S Parietooccipital 2 L | -13 | -65 | 26 | L | 12.735 |
| S Inf Frontal 2 L | -43 | 15 | 29 | L | 54.038 |

**Table S7.** List of significant regions for γ_H_ band (p < 0.01). Color code are Pink: Assortative; Green: Disassortative; Gray: Core-periphery.

| region | x | y | z | side | value [%] |
| --- | --- | --- | --- | --- | --- |
| G Occipital Pole 1 L | -21 | -96 | -14 | L | 45.216 |
| G Occipital Lat 1 L | -15 | -97 | 1 | L | 42.191 |
| G Occipital Lat 1 R | 17 | -94 | 4 | R | 43.067 |
| G Occipital Lat 2 L | -26 | -94 | -1 | L | 43.788 |
| G Occipital Lat 2 L | -40 | -84 | -12 | L | 43.699 |
| G Occipital Lat 2 R | 43 | -81 | -10 | R | 42.56 |
| G Occipital Lat 5 L | -35 | -79 | -1 | L | 43.44 |
| G Occipital Lat 5 R | 36 | -76 | 2 | R | 43.964 |
| G Occipital Mid 2 R | 41 | -73 | 12 | R | 43.678 |
| G Occipital Inf 1 R | 50 | -60 | -9 | R | 44.752 |
| G Occipital Inf 2 L | -45 | -71 | -7 | L | 42.12 |
| G Occipital Inf 2 R | 47 | -65 | -7 | R | 46.598 |
| G Temporal Mid 4 R | 57 | -53 | 3 | R | 44.371 |
| G Temporal Inf 3 R | 57 | -46 | -14 | R | 44.863 |
| G Temporal Inf 4 L | -50 | -61 | -8 | L | 43.253 |
| G Temporal Inf 4 R | 54 | -58 | -11 | R | 44.574 |
| G Temporal Inf 5 R | 49 | -58 | 4 | R | 45.088 |
| G Calcarine 3 R | 11 | -94 | 1 | R | 43.68 |
| G Lingual 2 R | 21 | -60 | -6 | R | 42.601 |
| G Lingual 4 L | -14 | -76 | -10 | L | 42.092 |
| G Lingual 5 L | -11 | -81 | -9 | L | 42.763 |
| G Fusiform 4 R | 44 | -46 | -18 | R | 43.543 |
| G Fusiform 6 R | 29 | -62 | -9 | R | 43.974 |
| G Fusiform 7 L | -23 | -84 | -10 | L | 44.402 |
| S Sup Frontal 5 R | 27 | 17 | 52 | R | 11.946 |
| G Supp Motor Area 1R | 6 | 21 | 48 | R | 13.37 |
| S Cingulate 2 R | 8 | 14 | 46 | R | 13.00 |
| S Cingulate 3 L | -7 | 5 | 47 | L | 12.66 |
| S Cingulate 3 R | 8 | 5 | 51 | R | 12.378 |
| G Cingulum Post 1 L | -4 | -26 | 29 | L | 12.001 |
| N Thalamus 2 L | -9 | -9 | 13 | L | 12.448 |
| N Thalamus 2 R | 9 | -7 | 13 | R | 12.619 |
| N Thalamus 3 L | -3 | -7 | -1 | L | 12.222 |
| N Thalamus 4 R | 3 | -14 | 9 | R | 12.345 |
| N Thalamus 5 R | 13 | -17 | 6 | R | 12.263 |
| N Thalamus 8 R | 4 | -25 | 13 | R | 12.25 |
| S Rolando 4 L | -23 | -29 | 64 | L | 60.739 |
| G Parietal Sup 2 L | -30 | -51 | 64 | L | 60.55 |

### Regions invariant to partitioning and showing the highest degree of community motif

The following tables are associated with the figure 5 of the manuscript. Indeed, they areas which are insensitive to portioning and showing a significant level of community motif. These show, frequency by frequency, the significant node’s name, MNI coordinate, hemisphere (according to the AICHA atlas).

**Table S8.** Regions in the δ band. Color code are Pink: Assortative; Green: Disassortative; Gray: Core-periphery.

| region | x | y | z | side | cluster |
| --- | --- | --- | --- | --- | --- |
| N Caudate 5 L | -13 | 10 | 8 | L | clust2 |
| G Precuneus 4 L | -7 | -49 | 45 | L | clust3 |
| G Occipital Inf 1 L | -48 | -69 | -4 | L | clust1 |
| G Temporal Mid 4 L | -53 | -59 | 7 | L | clust1 |
| G Temporal Inf 5 L | -45 | -64 | 6 | L | clust1 |
| G Temporal Sup 1 L | -55 | -1 | 2 | L | clust2 |
| G Temporal Mid 1 L | -61 | -12 | -17 | L | clust2 |
| G Temporal Inf 1 L | -47 | -15 | -33 | L | clust2 |
| G Occipital Pole 1 R | 24 | -93 | -13 | R | clust3 |
| G Occipital Lat 2 R | 28 | -89 | -2 | R | clust3 |
| G Occipital Lat 2 R | 43 | -81 | -10 | R | clust3 |
| G Occipital Lat 4 R | 34 | -85 | 9 | R | clust3 |
| G Occipital Lat 5 R | 36 | -76 | 2 | R | clust3 |
| G Occipital Mid 3 R | 45 | -63 | 15 | R | clust3 |
| G Occipital Inf 1 R | 50 | -60 | -9 | R | clust3 |
| S Sup Temporal 5 R | 51 | -54 | 21 | R | clust3 |
| G Temporal Inf 4 R | 54 | -58 | -11 | R | clust3 |
| G Lingual 5 R | 11 | -79 | -7 | R | clust3 |
| G Fusiform 7 R | 23 | -80 | -8 | R | clust3 |

**Table S9.**
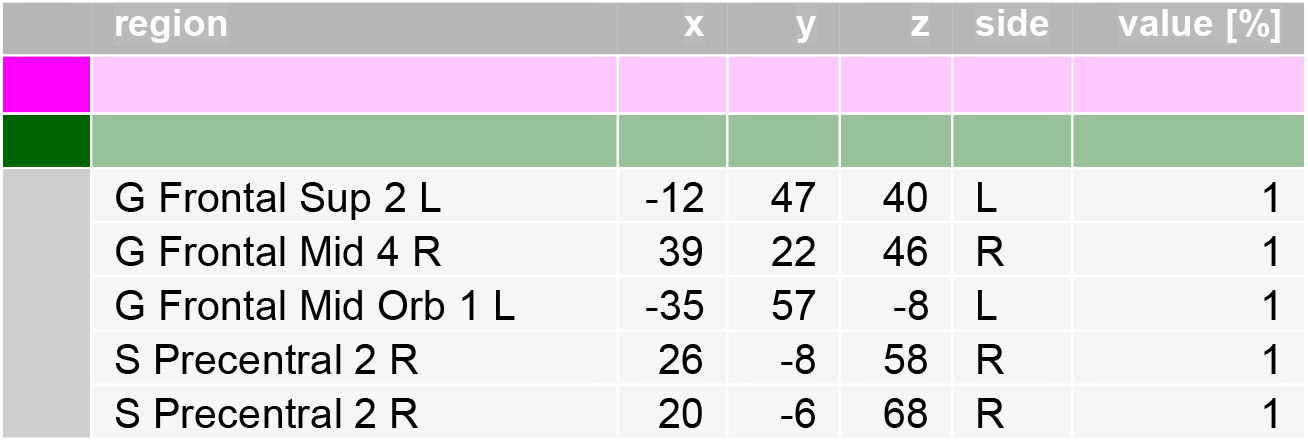
Regions in the θ band. Color code are Pink: Assortative; Green: Disassortative; Gray: Core-periphery.

**Table S10.** Regions in the α band. Color code are Pink: Assortative; Green: Disassortative; Gray: Core-periphery.

| region | x | y | z | side | cluster |
| --- | --- | --- | --- | --- | --- |
| G Frontal Mid 1 L | -40 | 41 | 20 | L | clust3 |
| S Inf Frontal 1 L | -44 | 38 | 12 | L | clust3 |
| G Frontal Inf Orb 1 L | -42 | 31 | -17 | L | clust3 |
| S Orbital 2 L | -31 | 34 | -13 | L | clust3 |
| G Frontal Sup Medial 1 L | -8 | 52 | 12 | L | clust3 |
| G Frontal Sup Medial 2 L | -7 | 53 | 25 | L | clust3 |
| S Anterior Rostral 1 L | -7 | 47 | -3 | L | clust3 |
| G Frontal Med Orb 2 L | -5 | 49 | -12 | L | clust3 |

**Table S11.** Regions in the β band. Color code are Pink: Assortative; Green: Disassortative; Gray: Core-periphery.

| region | x | y | z | side | cluster |
| --- | --- | --- | --- | --- | --- |
| G Frontal Sup 2 L | -12 | 47 | 40 | L | clust1 |
| S Sup Frontal 1 L | -22 | 61 | -8 | L | clust1 |
| S Sup Frontal 2 L | -27 | 56 | 1 | L | clust1 |
| S Sup Frontal 2 L | -26 | 43 | 29 | L | clust1 |
| G Frontal Mid 1 L | -40 | 41 | 20 | L | clust1 |
| G Frontal Mid Orb 1 L | -35 | 57 | -8 | L | clust1 |
| G Frontal Mid Orb 2 L | -41 | 49 | -5 | L | clust1 |
| G Parietal Sup 2 L | -30 | -51 | 64 | L | clust2 |
| G Angular 2 R | 45 | -62 | 36 | R | clust2 |
| S Intraparietal 2 R | 37 | -52 | 48 | R | clust2 |
| S Cingulate 6 R | 11 | -34 | 46 | R | clust2 |
| S Cingulate 7 R | 9 | -41 | 61 | R | clust2 |
| G Cingulum Post 2 L | -4 | -39 | 27 | L | clust2 |
| G Cingulum Post 2 R | 8 | -43 | 31 | R | clust2 |
| G Precuneus 3 L | -7 | -51 | 34 | L | clust2 |
| G Precuneus 3 R | 6 | -52 | 32 | R | clust2 |
| G Precuneus 4 L | -7 | -49 | 45 | L | clust2 |
| G Precuneus 4 R | 8 | -48 | 43 | R | clust2 |
| G Precuneus 5 L | -8 | -55 | 55 | L | clust2 |
| G Precuneus 5 R | 7 | -50 | 53 | R | clust2 |
| N Thalamus 7 R | 9 | -26 | 10 | R | clust2 |
| G Frontal Inf Tri 1 R | 50 | 29 | 5 | R | clust1 |
| G Frontal Mid Orb 2 R | 40 | 50 | -4 | R | clust1 |

**Table S12.** Regions in the γ_L_ band. Color code are Pink: Assortative; Green: Disassortative; Gray: Core-periphery.

| region | x | y | z | side | cluster |
| --- | --- | --- | --- | --- | --- |
| G Occipital Pole 1 R | 24 | -93 | -13 | R | clust1 |
| G Occipital Lat 1 L | -15 | -97 | 1 | L | clust1 |
| G Occipital Lat 1 R | 17 | -94 | 4 | R | clust1 |
| G Occipital Lat 2 R | 28 | -89 | -2 | R | clust1 |
| G Occipital Lat 2 R | 43 | -81 | -10 | R | clust1 |
| G Occipital Lat 4 R | 34 | -85 | 9 | R | clust1 |
| G Occipital Lat 5 R | 36 | -76 | 2 | R | clust1 |
| G Occipital Sup 1 R | 18 | -81 | 37 | R | clust1 |
| G Occipital Sup 2 R | 17 | -88 | 26 | R | clust1 |
| G Occipital Mid 2 R | 41 | -73 | 12 | R | clust1 |
| G Occipital Inf 1 L | -48 | -69 | -4 | L | clust1 |
| G Occipital Inf 1 R | 50 | -60 | -9 | R | clust1 |
| G Occipital Inf 2 L | -45 | -71 | -7 | L | clust1 |
| G Occipital Inf 2 R | 47 | -65 | -7 | R | clust1 |
| G Temporal Inf 4 L | -50 | -61 | -8 | L | clust1 |
| G Temporal Inf 4 R | 54 | -58 | -11 | R | clust1 |
| G Cuneus 2 L | -4 | -92 | 19 | L | clust1 |
| G Cuneus 2 R | 8 | -89 | 19 | R | clust1 |
| G Calcarine 3 L | -7 | -95 | -1 | L | clust1 |
| G Calcarine 3 R | 11 | -94 | 1 | R | clust1 |
| G Lingual 4 R | 13 | -72 | -9 | R | clust1 |
| G Lingual 5 R | 11 | -79 | -7 | R | clust1 |
| G Lingual 6 R | 7 | -79 | -3 | R | clust1 |
| G Fusiform 7 R | 23 | -80 | -8 | R | clust1 |

**Table S13.**
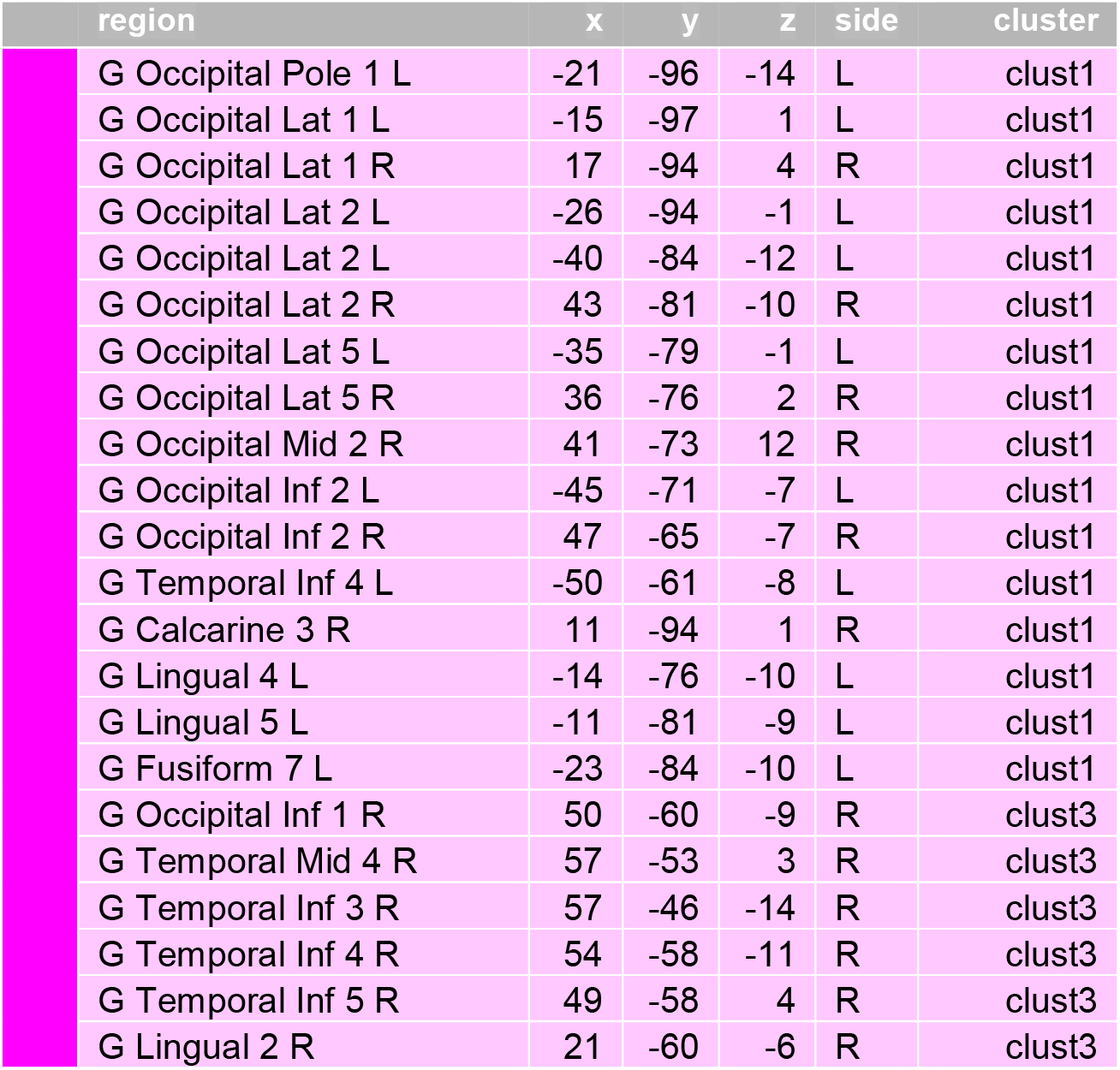
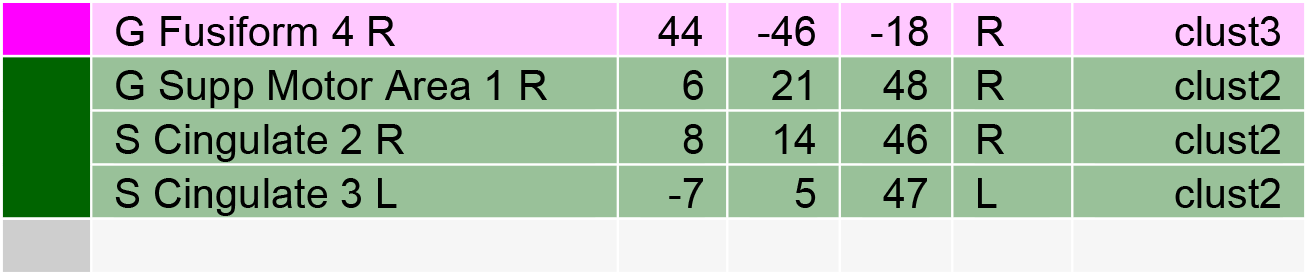
Regions in the γ_H_ band. Color code are Pink: Assortative; Green: Disassortative; Gray: Core-periphery.

